# Single Cell Variability of CRISPR-Cas Interference and Adaptation

**DOI:** 10.1101/2021.07.21.453200

**Authors:** Rebecca E. McKenzie, Emma M. Keizer, Jochem N.A. Vink, Jasper van Lopik, Ferhat Büke, Vera Kalkman, Christian Fleck, Sander J. Tans, Stan J.J. Brouns

## Abstract

CRISPR-Cas defence is a combination of adaptation to new invaders by spacer acquisition, and interference by targeted nuclease activity. While these processes have been studied on a population level, the individual cellular variability has remained unknown. Here, using a microfluidic device combined with time-lapse microscopy, we monitor invader clearance in a population of *Escherichia coli* across multiple generations. We observed that CRISPR interference is fast with a narrow distribution of clearance times. In contrast, for invaders with escaping PAM mutations we show large cell-to-cell variability of clearance times, which originates from primed CRISPR adaptation. Faster growth and cell division, as well as higher levels of Cascade, increase the chance of clearance by interference. In contrast, faster growth is associated with decreased chances of clearance by priming. A mathematical model explains the experimental findings, and identifies Cascade binding to the mutated invader DNA, rather than spacer integration, as the main source of priming heterogeneity. The highly stochastic nature of primed CRISPR adaptation implies that only subpopulations of bacteria are able to respond to invading threats in a timely manner. We conjecture that CRISPR-Cas dynamics and heterogeneity at the cellular level are crucial to understanding the strategy of bacteria in their competition with other species and phages.

## Introduction

During the last decade, important progress has been made in identifying the sequence of steps and molecular interactions required for successful adaptive immunity by the model type I-E CRISPR-Cas system ^1–10^. CRISPR (Clustered Regularly Interspaced Short Palindromic Repeats) immunity involves three main stages beginning with the acquisition of a spacer, a small piece of DNA derived from a foreign invader and stored in the CRISPR array for future defence ^11, 12^. This array is then transcribed and processed into small CRISPR RNAs (crRNAs) which guide a surveillance complex, formed from a number of Cas (CRISPR-associated) proteins, towards the invaders DNA ^13, 14^. For type I-E systems a 5’-CTT consensus PAM (Protospacer Adjacent Motif) sequence flanking the targeted site of the invader ^15, 16^ is required for recognition and ultimately degradation of the invader, through a process called direct interference ^7, 17–19^. However, invaders can escape direct interference via mutation within the seed region of the target site or PAM ^15, 20, 21^. In response, the I-E system can initiate priming, which promotes accelerated acquisition of new spacers due to a pre-existing partial match to the invader ^1, 4^. Primed adaptation is much faster than naïve adaptation ^22^, and is required for the insertion of a new matching spacer with a consensus PAM allowing subsequent invader degradation, which we here refer to as primed interference.

At the level of individual cells however, much more is unknown. Interference is a tug-of-war between invader replication and degradation, which could result in complex and stochastic dynamics within single cells. Replication and degradation themselves may also display variability between cells in the population. For instance, invader degradation rates can be affected by stochastic processes such as the expression of CRISPR-Cas components, target localization, and nuclease recruitment ^3, 23^. Priming also depends on many processes in which the dynamical interplay is unclear, including the production of suitable fragments of DNA for spacer acquisition (pre-spacers), the assembly of adaptation complexes required for further spacer selection, and the processing and insertion of these pre-spacers into the CRISPR array ^6, 10, 24, 25^. Elucidating the cellular dynamics and heterogeneity of the CRISPR-Cas response is critical to understanding interference and adaptation mechanistically, and of direct importance to its natural function. For instance, upon invasion, cells are thought to have a limited time window to respond in order to escape invader replication, protein production, and cell death ^26–29^.

A number of studies have investigated the interference process by collecting either population averages, or single-cell data on short time scales (<1 s) ^3, 5, 21, 30–32^. However, averaging within a population can conceal the variation between cells, as well as the dynamics within single-cells over time ^33, 34^, thus masking the underlying dynamics of CRISPR-Cas interference. In addition, investigations into the adaptation process have provided great insight into the diversity of spacers acquired ^30, 35^, possible mechanisms of target destruction ^1, 36^, and conditions under which adaptation most frequently occurs within a population ^37–39^, however these studies could not observe any variation existing in each step of the adaptation process within individual cells.

Here we set out to investigate and quantify the dynamics and variability of the CRISPR-Cas response at the single-cell level. Using time-lapse microscopy and microfluidic devices, we followed individual cells over multiple rounds of division while simultaneously monitoring CRISPR-Cas protein expression and DNA degradation. Hence, we obtained individual lineages, the genealogical relations between them, as well as real-time data on the DNA clearance process, instantaneous growth rates, cell sizes, and division frequencies of individual cells. We determined that while direct interference occurs quickly and consistently, clearing the target from all cells within hours, priming is highly variable and much slower, taking up to several tens of hours. Further, we were able to define the adaptation and clearance stages of priming and identified primed adaptation as the source of the variation observed. Finally, we corroborated our findings with a minimal agent-based model, that accurately replicated our data and provided further insights into the dynamics of the primed adaptation process.

## Results

### Time-lapse microscopy of the CRISPR-Cas response

Using two strains, KD615 (WT) and KD635 (Δ*cas*1,2) (Supplementary Table 1), we investigated priming and direct interference respectively. The strains contain an array with a leader, two repeats and a single previously characterized spacer, spacer8 (SP8) ^4, 5^ (Fig. 1a-c). In addition, these strains are engineered to control *cas* gene expression using arabinose and IPTG induction, and hence initiation of the CRISPR-Cas response. Target plasmids were engineered to encode a constitutively expressed YFP or CFP fluorescent protein ^40^ and contain a target sequence that is complementarity to SP8 in the CRISPR array, allowing direct monitoring of target DNA presence in individual cells over time (Fig. 1a-c) (Supplementary Table 1). In order to investigate the direct interference process, we flanked the target sequence with a 5’-CTT consensus PAM ^16^ (Fig 1a,b). Further, to investigate the priming response we mutated the PAM to 5’-CGT (Fig 1b,c), a mutation known to allow mobile genetic elements (MGE) to escape interference, and invoke a primed adaptation response ^1, 5, 20^.

**Fig. 1.**
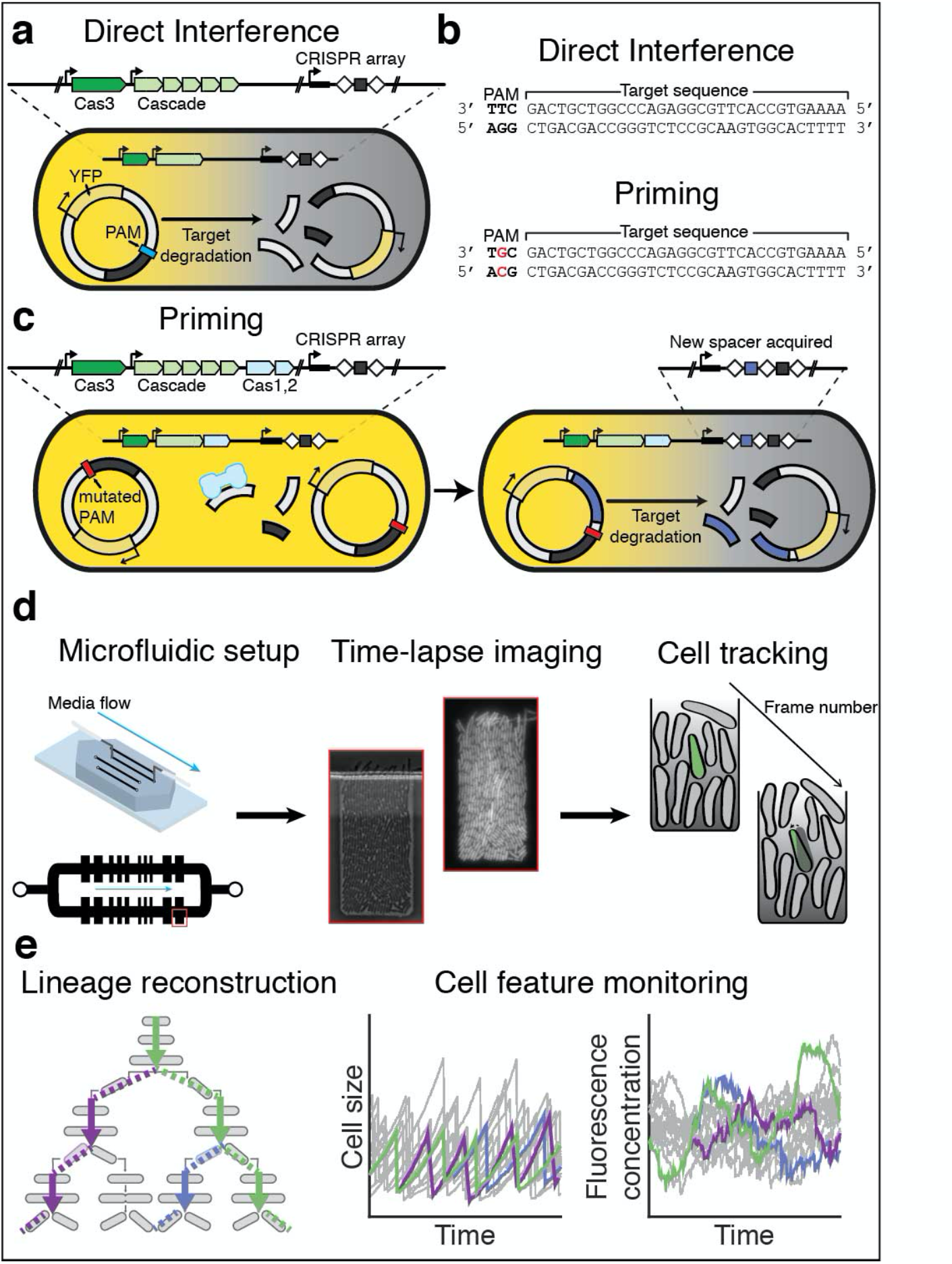
Investigating single-cell behaviour during CRISPR-Cas defence using time-lapse microscopy. **a,** Schematic of the direct interference process. The cell contains a I-E CRISPR-Cas system, as well as the CRISPR array with a single spacer targeting the plasmid (grey box). The plasmid encodes YFP and contains a sequence matching the spacer (grey), flanked by a consensus PAM (blue). Immediate targeting by the CRISPR-Cas system resulting in degradation of the plasmid and loss of the YFP in the cell. **b,** To invoke priming the 5’-CTT consensus PAM, flanking the target sequence located on the plasmid, is mutated by one nucleotide to a non- consensus PAM 5’-CGT. **c,** Schematic of the priming process. (Left cell) A mutation of the PAM (red) flanking the target sequence means the spacer in the CRISPR array can no longer initiate direct interference. Fragments in the cell can be captured and processed by Cas1,2 (light blue). (Right cell) The Cas1,2 complex integrates the fragment into the CRISPR array as a new spacer (purple), which matches the target plasmid resulting in degradation and loss of YFP in the cell. **d,** To allow long term imaging cells are grown in a microfluidic chip that allows constant media supply. Cells within a single well are imaged frequently in phase contrast and fluorescence allowing segmentation and tracking of lineage history across frames. **e,** Variation in features of reconstructed single-cell lineages (left) such as size (middle) and fluorescence concentration (right) are continuously monitored enabling further investigation.

Use of a microfluidic device ^41^ enabled fluorescence time-lapse imaging for over 36 hours with the option for media exchange (Fig. 1d). The device contained chambers allowing observation of a single layer of cells, constant medium supply, removal of cells that no longer fit the chamber due to growth, and control of intracellular processes via induction. Image analysis software was used to segment and track all cells and their fluorescence signals, thus allowing the re-construction of lineage trees in a defined region at the bottom of the chamber (Fig. 1d,e)^41–43^.

### Direct interference is fast and synchronous

We first investigated the direct interference response (Fig. 1a). Prior to *cas* gene induction, the images showed high YFP fluorescence in all cells, confirming the presence of the target plasmid (Fig. 2a) which decreased upon induction, indicating CRISPR-Cas mediated degradation of the target DNA (Fig. 2a, Supplementary video 1). When the plasmid did not contain a target sequence (pControl) YFP levels did not decrease for over 35 hours (Supplementary Fig. 1), indicating that the plasmid loss was CRISPR-Cas dependent.

**Fig. 2.**
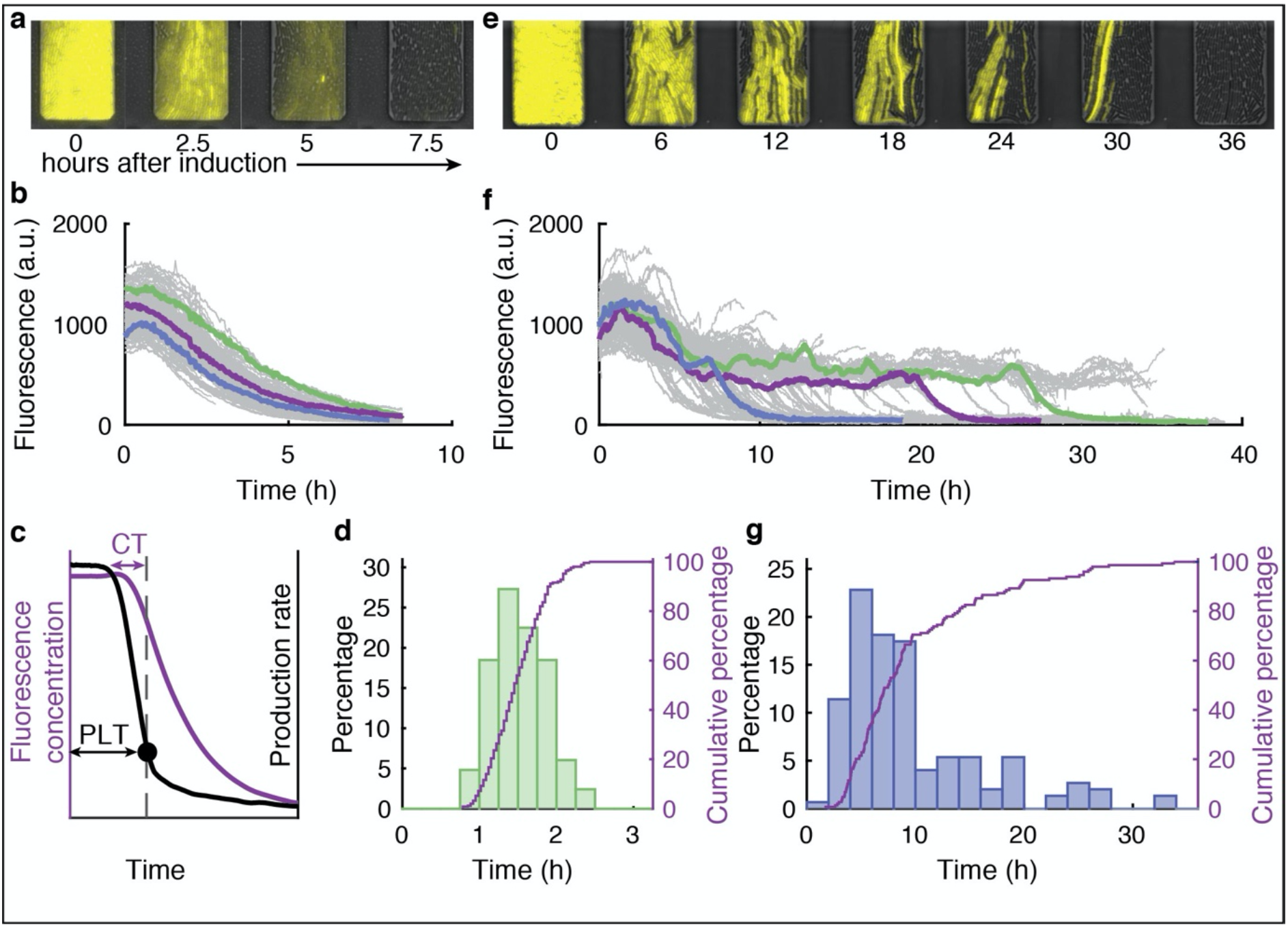
Variation in target plasmid clearance times is much larger when CRISPR adaptation is required. **a-b**, Depict clearance of a target with a consensus PAM by direct interference **a,** Overlay of fluorescent and phase contrast time-lapse images. Presence of the target plasmid is tracked by its YFP production. Images are shown at 2.5 h intervals starting from induction of *cas* gene expression. **b**, Reconstructed lineage traces of the imaged population (a) from induction of the CRISPR-Cas system over time (grey) lineages show some variation in plasmid clearance times (coloured). **c**, Production rate (black line) of the YFP is used to determine the plasmid loss time, PLT, (black dot, black arrow) allowing earlier detection than using the mean fluorescence (purple line). The time from first targeting of a single plasmid to the PLT is defined as the clearance time (CT, purple arrow). **d**, Distribution of PLTs determined by the production rate during direct interference (n=250). **e-f**, Depict clearance of a matching target with a mutated PAM via priming **e,** Overlay of fluorescent and phase contrast time-lapse images. Presence of the target is tracked by YFP production. Images are shown at 6 h intervals. **f**, Reconstructed lineage traces of the imaged population (e) from induction of the CRISPR-Cas system over time (grey). Lineages show large variations in the time taken to clear the plasmid (coloured). **g**, Distribution of plasmid loss times calculated with the production rate during priming (n=149).

The mean YFP fluorescence per cell unit area (which estimates the YFP concentration) showed the decrease started after about 1 hour of induction, and then exhibited a smooth monotonic decline without significant fluctuations (Fig. 2b). Note that traces end upon the cells exiting the observation chamber. CRISPR mediated degradation of the target was thus efficient and synchronous, and in the case of a 5-copy plasmid could overcome the plasmid replication and copy number control. Hence, we surmised that the YFP fluorescence may decrease exponentially, as the YFP proteins are diluted exponentially due to volume growth upon clearance of the last plasmid. Indeed, we found the fluorescence decrease to be exponential (Supplementary Fig. 2).

Direct interference variability between cells also appeared limited (Fig. 2b). To address it more directly, we quantified the moment all plasmids are cleared by determining the YFP production rate as the change in total cellular fluorescence per unit of time ^44^. The production rate scales with the number of target DNA copies, and shows the expression timing more precisely by suppressing slow dilution effects. Indeed, the YFP production rate decreased rapidly, and reached zero (the background level of cells not expressing YFP) when the mean fluorescence was still close to its maximum (Fig. 2c). This moment was identified as the plasmid loss time (PLT) (Fig. 2c). PLT was narrowly distributed between about 1 and 2.5 hours (Fig. 2d, CV = 0.055). Hence, in all cells the target was cleared. The clearance was rapid taking between 1 and 3 generations, and sometimes occurred in the same generation in which the CRISPR-Cas response was initiated by induction (Supplementary Fig. 3).

### Primed adaptation is highly variable

Next, we studied plasmid clearance after adaptation from a target with a mutated PAM (Fig. 1c). Most notable in these priming experiments was the heterogeneity between lineages, with the clearance process ranging from 2-30 cellular generations (Supplementary Fig. 3). Upon induction, some lineages showed a decreasing trend in fluorescence as early as 4 hours (Fig. 2e-f, Supplementary video 2), while others remained fluorescent after 35 hours (Fig. 2f). The PLT’s were indeed broadly distributed and displayed a long tail towards large values (Fig. 2g, CV 0.458). Of note, we did not observe plasmid clearance in the same generation in which the CRISPR-Cas system was induced (Supplementary Fig. 3).

The shapes of the YFP declines were exponential, similar to the direct interference data (Fig. 2b and f, Supplementary Fig. 2). Alignment of all production rate traces at the PLT yielded a similar averaged profile for direct interference and priming (Supplementary Fig. 4). In these data, the onset of the decrease is about 60 min before PLT in both cases, thus estimating the clearance time (CT), the duration of the target clearance process. In priming, clearance thus contributes much less to PLT variability than the preceding processes (Fig. 2g). These observations suggest that new spacers preceded by a consensus PAM are indeed acquired, and that the CRISPR adaptation phase is responsible for the observed temporal variability (Fig. 2g).

Spacer acquisition in the population was indeed confirmed by PCR of the CRISPR array in cells collected from the microfluidic device (Supplementary Fig. 5). Spacer acquisition was not observed with the Δ*cas*1,2 strain, consistent with Cas1 and Cas2 being required for acquisition ^45^. In the absence of Cas1 and Cas2 however, low frequency plasmid loss was observed in 1.4% of the lineages over a 35-hour period (Supplementary Fig. 6). Hence, complete clearance is possible with a mutated PAM, even if highly inefficient.

### Genealogical relations impact the CRISPR-Cas response

To study the role of genealogy in the CRISPR-Cas response, we took a more in depth look at the lineage history before plasmid loss (Fig. 3a). For primed adaptation, some subtrees showed all plasmid loss events occurring close together, however most subtrees showed a wide PLT variability (Fig. 3b, black dots), in line with lineages responding independently. However, statistical analysis showed that sisters cleared their plasmids within the same cell cycle more frequently than expected at random, and more strongly so for priming than for direct interference (Fig. 3c). Hence, inheritance plays a role in the CRISPR-Cas response (Fig. 3c).

**Fig. 3.**
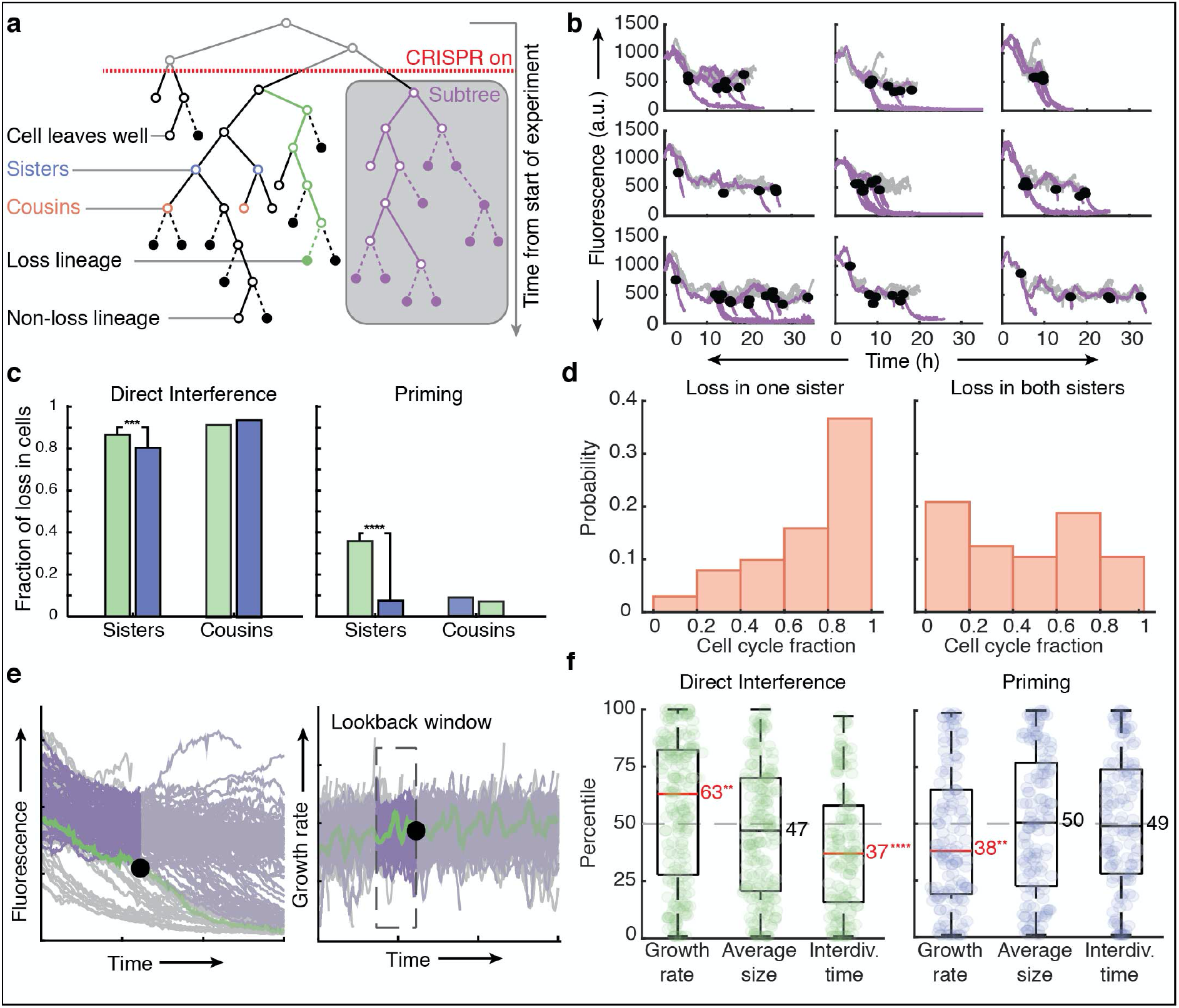
Growth rate and interdivision times have an influence on direct interference and priming. **a**, Schematic of key analysis structure and terminology. **b**, A comparison of 9 subtrees constructed from induction. **c**, The observed fraction of loss in cells (green) during direct interference (left) or priming (right) related as either sisters (DI: n=171, P: n=98) or cousins (DI: n=130, P: n=138) is plotted against the fraction of expected loss events (blue) in related cells when the events are randomized in the same time window. **d**, The cell cycle was divided into 5 equal fractions and plasmid loss times are plotted in the corresponding fraction where one sister alone cleared the plasmid (left, n=101) or both sisters cleared the plasmid (right, n=24) **e**, Schematic explaining the rank-based analysis approach. For each detected loss event (left, black circle) the cell feature i.e. growth rate for that lineage (right, green) is averaged over a lookback window (right, dashed rectangle), and then ranked amongst all averages of non-loss lineages in the same window (violet, right). **f**, Boxplots of percentile rankings of all loss lineages that cleared a consensus target via direct interference (green, left, n=250), or a mutated target via priming (blue, right, n=149), for growth rate, birth size and interdivision respectively over a lookback window of 30 minutes. The median percentile ranking of loss lineages is indicated by a line and value, categories in which this value was significantly different from a ranking in the 50^th^ percentile as computed by a 2-sided binomial test are indicated in red followed by asterisks. (****p<0.0001, ***p<0.001, **p<0.01)

These data led us to hypothesize that in priming, sisters correlate because spacer acquisition occurs in the mother, after which plasmid degradation (primed interference) continues into the daughters. If true, the moment of plasmid loss may be distributed throughout the daughter’s cell cycle, depending on the acquisition moment within the mother’s cell cycle. Conversely, when acquisition and clearance both manage to occur in the mother, hence yielding no correlated loss events, clearance should occur at the end of the mother’s cell cycle, because of the time (∼ 60 min) needed for plasmid clearance. To test this hypothesis, we divided the cell cycles into five fractions, and tabulated the observed loss event for each fraction. Indeed, loss events without correlated plasmid loss in the sister cell occurred more frequently towards the end of the cell cycle (Fig 3d), while the moment of loss was more randomly distributed when both sisters lost the plasmid (Fig 3d). Altogether this indicated that loss likely takes place more frequently in sisters than cousins (Fig 3c) because adaptation occurred in the mother.

### The growth rate has opposing effects on adaptation and interference

To study if stochastic variations in cell cycle parameters affect the CRISPR-Cas response, we used a ranking analysis, as this suppresses long-term trends in the population (Fig. 3e). For instance, we ranked each lineage showing plasmid loss relative to lineages that had not lost their plasmids at that moment, with the ranking based on growth rate averaged over a ‘lookback window’ (Fig. 3e), determined using autocorrelation times (Supplementary Fig. 7). In direct interference, the ‘loss lineages’ exhibited a higher median growth rate than ‘non-loss lineages’, with their growth rate ranking in the in the 63^rd^ percentile (p=0.01) (Fig. 3f). These lineages also showed shorter interdivision times (p=0.0001), but not a difference in cell size (Fig. 3f). These results were robust over a range of lookback window sizes (see Supplementary Fig. 8). We stress that growth is likely only one of the many factors affecting the CRISPR-Cas response, which is also reflected by the broad ranking distributions (Fig. 3f). Overall, the analysis indicated that faster growth in coordination with more frequent cell division has a positive effect on the rate of clearance of a consensus target.

Primed adaptation showed a different picture. To probe the effects on spacer acquisition, which occur about 60 min before plasmid loss, we used a lookback window between 90 and 60 min before the PLT. While cell size and interdivision time did not show an effect (no significant deviation from the 50^th^ percentile) the growth rate did, with loss lineages growing more slowly compared to non-loss lineages (38^th^ percentile, p=0.01) (Fig. 3f). This was robust to changes in the lookback window (Supplementary Fig. 9). Altogether, these findings indicated that, on average, slower growing cells achieved faster plasmid clearance through priming.

### Cascade concentrations impact the CRISPR-Cas response

Apart from physiological determinants like growth ^46^, Cascade expression levels may influence the speed of CRISPR-Cas defence. We fused mCherry (RFP) to the N-terminus of the Cas8e subunit of Cascade ^3^ (Fig. 4a). Using single particle fluorescence calibration, we estimated that the cells contain on average about 200 Cascade molecules/µm^2^ (Fig. 4b and Supplementary Fig. 10). Hence, we quantified the (stochastic) variations in Cascade abundance within single-cell-lineages upon induction (Fig. 4b).

**Fig. 4.**
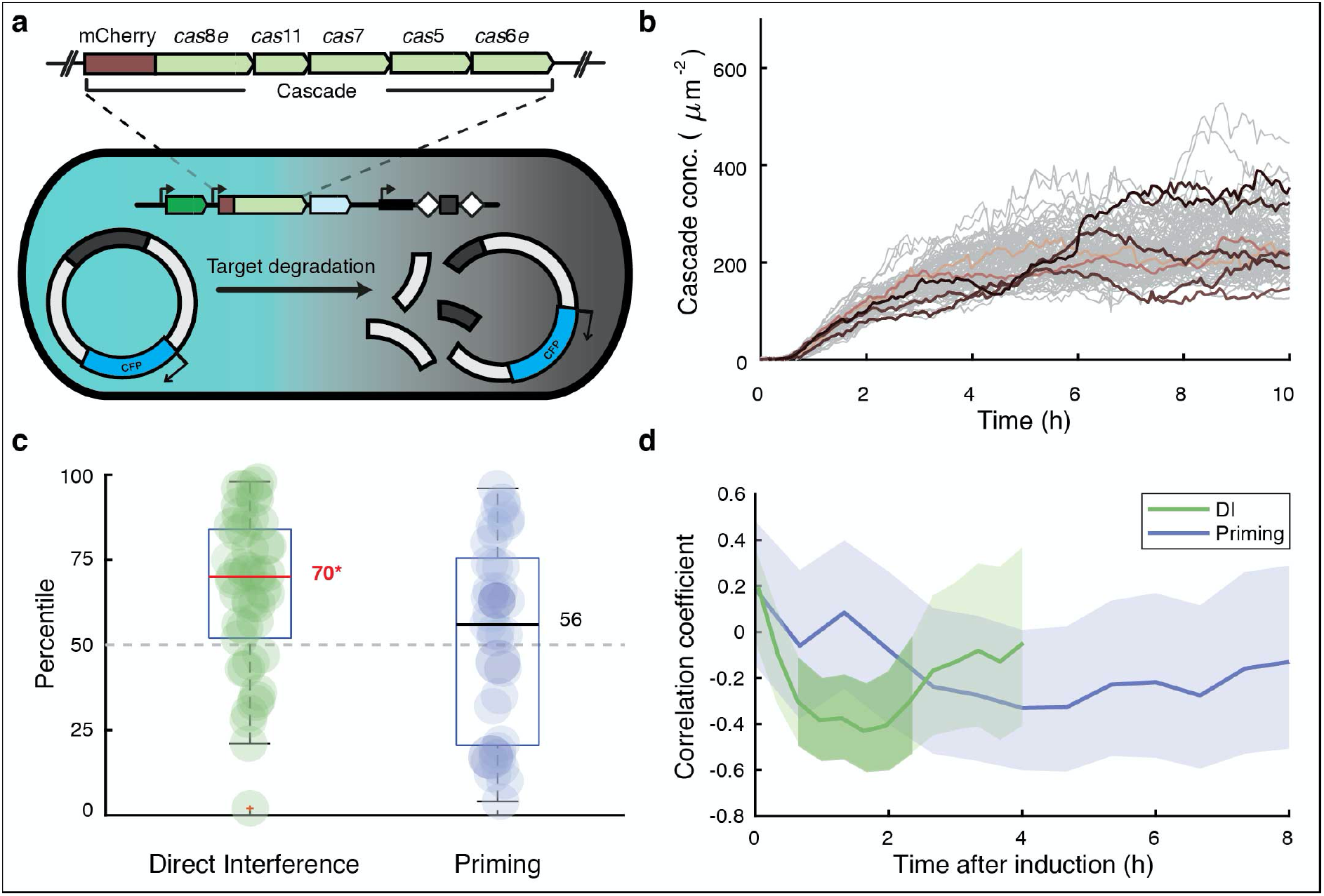
Growth rate and interdivision times influence direct interference and priming. **a**, Schematic of the experimental set up adapted to allow visualization of target presence (CFP) and Cascade levels (mCherry) simultaneously. The expansion indicates the mCherry fluorescent tag was integrated upstream of the *cas8e* subunit. **b**, Cascade concentration of single-cell lineages over time from induction. **c**, Cascade concentrations were averaged over a 30-minute lookback window from the plasmid loss event for all loss lineages during direct interference (green) or priming (blue). The Cascade concentration of the loss lineages were ranked as percentile amongst the non-loss lineages and plotted here. The median percentile ranking of loss lineages is indicated by a line and value, categories in which this value was significantly different from a ranking in the 50^th^ percentile as computed by a 2-sided binomial test (*p<0.05) are indicated in red followed by an asterisk. **d**, The Pearson correlation coefficient of plasmid loss time versus total cumulative Cascade concentration at that moment is plotted every 5 minutes (DI) or 10 minutes (Priming) starting from induction of the CRISPR-Cas system. The plotted line for both a target with a consensus PAM (green) and target with a mutant PAM (blue) are enveloped by a 95% confidence interval. Darker shading indicates where the correlation coefficient is significantly different from zero (p<0.05).

Cascade levels fluctuated on a longer timescale than the cell cycle (200 min, Supplementary Fig. 11) and were strongly correlated between sisters and cousins (R=0.89 and 0.62 respectively, Supplementary Fig. 12) indicating that Cascade levels are stable over several generations. We reasoned that lineages with high Cascade concentrations may target and clear the plasmids faster. Hence, we performed time-lapse experiments and used the ranking approach, now ranking lineages based on the average Cascade in a window 60 min prior to plasmid loss. For direct interference, loss lineages exhibited significantly higher Cascade levels than non-loss lineages, and ranked in the 70^th^ percentile (p=0.03, Fig. 4c). Conversely, no differences in Cascade levels were observed between loss and non-loss lineages for priming, with the former ranking in the 56^th^ percentile (Fig. 4c).

In priming however, the target search by Cascade occurs over a longer period of time prior to achieving plasmid loss, likely rendering the ranking approach less suitable. Hence, to test this notion, we quantified the total area under the Cascade concentration curves (Fig. 4b) in a certain time window. At 0-2 hours post induction, PLT and Cascade search hours indeed correlated negatively for direct interference but not for priming (Fig. 4d), in line with our earlier analysis (Fig. 4c). Interestingly, beyond 2 hours post induction, the correlation increases in magnitude for priming, even as the variability is high, and the correlation is not significant (Fig. 4d). In summary, stochastic variations in Cascade expression levels affect direct interference. For the priming process, the impact of Cascade levels appeared weaker. We hypothesize this could be due to the underlying processes being less synchronized in time in comparison to direct interference, and hence masked by other stochastic variations.

### Low Cascade-target binding affinity generates CRISPR-Cas response variability

To gain insight into the variability and dynamics of the CRISPR-Cas defence we developed an agent-based simulation framework. Adaptive immunity in bacterial populations has been modelled previously ^47–49^ but to our knowledge none describe variability or single-cell behaviour. Briefly, we simulated 100 cells, their growth and division, plasmid maintenance, stochastic protein production and partitioning at division, spacer acquisition, and target DNA degradation (see Supplementary Methods for details). We found that with these minimal model ingredients, and by only changing the Cascade-target binding affinity due to the PAM mutation, the model could reproduce both the low variability of direct interference (Fig. 5a,b and Fig. 2b,d), and the high variability of priming (Fig. 5c,d and Fig. 2f,g) from the experimental conditions. Specifically, we found a Cascade-target binding affinity reduction of two orders of magnitude for the PAM mutation, which is consistent with previous work ^50, 51^ (Supplementary Methods Table 2).

**Fig. 5.**
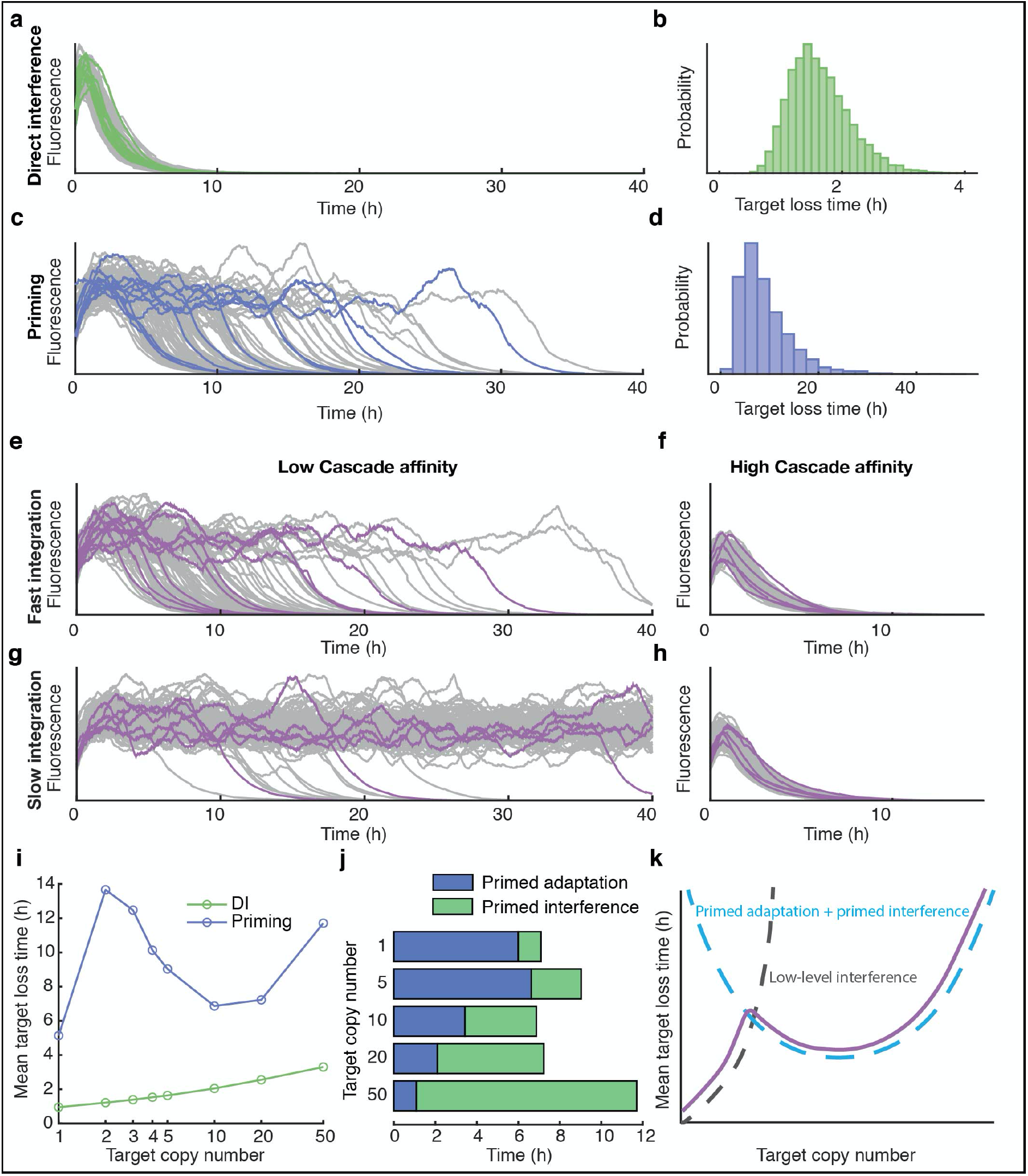
Results from the stochastic agent-based model of CRISPR adaptation and interference. **a-d**, Example trajectories showing fluorescence concentration produced by target plasmids simulated with the agent-based model for the (**a**) direct interference and (**c**) priming condition, and corresponding target loss distribution (**b,d** respectively). **e-h**, Example trajectories from 4 different parameter combinations. High Cascade affinity (**f,h**) corresponds an increase in target binding by a factor 100 as compared to low Cascade affinity (**e,g**), slow integration (**g,h**) represents a 100-fold reduction in the spacer integration rate as compared to fast integration (**e,f**). **i**, Mean target loss time of the population as a function of the average target copy number per cell for direct interference (green) and priming (blue). **j,** Breakdown of average time spent on primed adaptation (blue) and primed interference (green) for cells that clear targets through priming, for target copy numbers in the range 1-50. **k**, Schematic of alternative target loss pathways. At low copy numbers, targets can be completely cleared through low-level interference, which becomes increasingly rare as copy numbers increase. The priming process shows a u-shaped relationship with the target copy number, as a result of adaptation becoming faster as target copy numbers increase, and time required for interference increasing with target copy number.

The priming process can be conceptually understood as a two-step process, adaptation followed by interference, where a highly reduced rate of the first step is able to recreate the broadness of the PLT distribution (see Supplementary Methods for details). We hypothesized that variation of the primed adaptation response could originate from the low-affinity target search of Cascade, or the spacer integration. In the agent-based model, we were able to vary the rates of these two processes by a factor 100, while keeping the Cas3- mediated target destruction constant, and find that slow spacer integration alone is not enough to explain the observed variability (Fig. 5h). Conversely, reduced Cascade-target binding affinity is both necessary and sufficient to reproduce the observations (Fig. 5e-h) and is required to generate pre-spacers.

### Competition between adaptation and low-level interference

In priming, low Cascade-target affinity and resulting sporadic target degradation can yield a low-level interference prior to adaptation, which in turn provides a continuous source of target DNA fragments that can act as pre-spacers ^2^. Hence, we wondered whether target abundance affects this process. For direct interference, as expected, we found that the PLT increased monotonically in simulated trajectories as the average number of targets varies from 1 to 50 (Fig. 5i, see Supplementary Fig. 13 for full range of distributions). Simulations of priming did not show such a monotonic trend: the PLT first went up, then down, and finally up again (Fig. 5i, Supplementary Fig. 14). This behaviour could be explained by splitting priming into adaptation and interference (Fig. 5j): while primed interference logically only speeds up with fewer targets, primed adaptation initially slows down with fewer targets because of the resulting fewer pre-spacers, but then speeds up for the lowest number of targets, because low-level interference is now sufficiently efficient, in combination with unequal partitioning upon division (Supplementary Fig. 15). Indeed, our experiments also showed such clearance of a 5-copy target by low-level interference without spacer acquisition (Supplementary Fig. 3). This alternative pathway competes with priming when there are few targets (Fig. 5k), and might explain the trend in Fig. 5j showing faster loss at 1 target as compared to 5 targets. Target abundance thus affects the balance between primed adaptation and primed interference, resulting in a non-monotonous trend for the target clearance probability.

### Cascade expression stochasticity can accelerate CRISPR adaptation

Our experiments showed that CRISPR-Cas defence is affected by Cascade expression (Fig. 4c-d) which is stochastic in nature (Fig. 4b). For direct interference simulations, we changed the level of gene expression variability for Cascade to have 100-fold stronger expression bursts while maintaining average Cascade concentrations (see Supplementary Methods for details), which resulted in a higher mean PLT: while some cells could clear all targets earlier, many cells required more time to clear all targets as compared to lower-variability Cascade expression (Supplementary Fig. 16). Surprisingly, for priming the mean PLT became lower when the Cascade variability increased (Fig. 6a). The primed interference phase showed a trend similar to direct interference: a broadening of the PLT distribution yielding a slow-down on average (Fig. 6b). However, the entire distribution shifted to lower values for primed adaptation (Fig. 6c), yielding an overall speed-up. For mutated PAMs, pre-spacer production critically depends on high Cascade levels, even if transient, as the cumulative probability of a pre-spacer integration event depends on the Cas concentration in a highly non-linear fashion (see Supplementary Methods for a more detailed illustration).

**Fig. 6.**
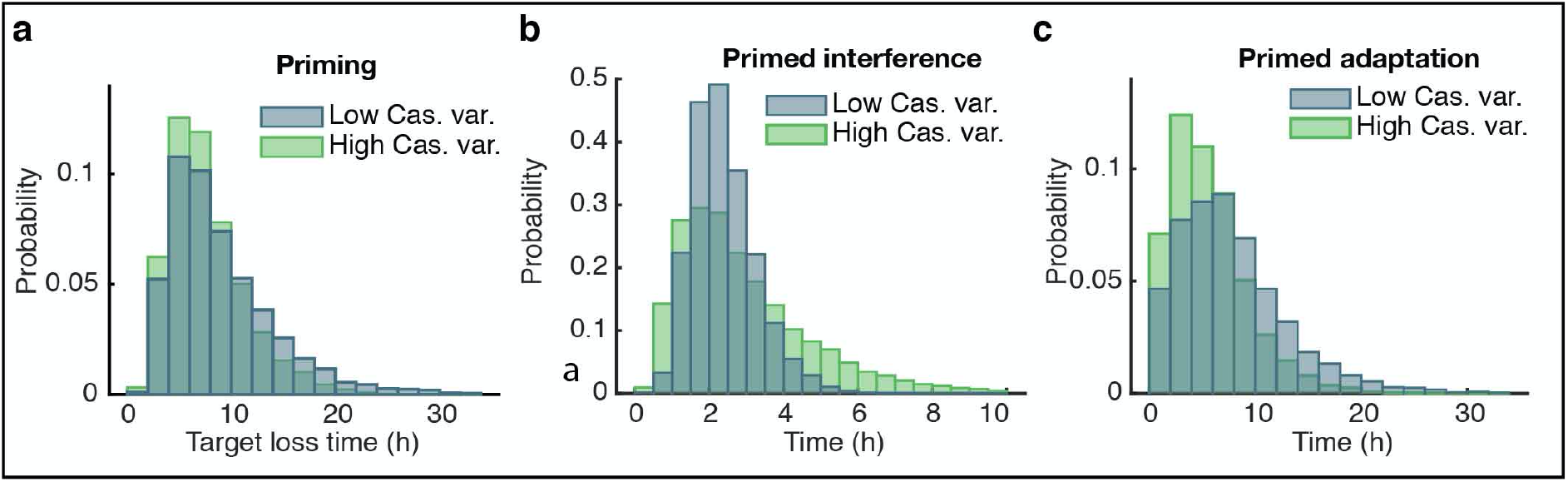
Distribution of primed adaptation and primed interference time for high and low variability in Cascade concentration. **a,** Target loss time distribution for two different levels of Cascade concentration variability for priming. At low variability (blue) Cascade proteins are produced in frequent, small bursts, whereas at high variability (green) proteins are synthesized more sporadically in large bursts (100-fold increase), keeping average Cascade concentration constant. **b-c,** The variability of primed interference times (**b**) for high Cascade variability (green) increases as compared to low Cascade variability (blue), whereas the variability of primed adaptation times (**c**) decreases with higher Cascade variability.

## Discussion

In this study, we have investigated a previously unexplored question: what are the dynamics and variability of the CRISPR adaptation and interference responses in individual cells? Our time-lapse microscopy approach allowed real-time monitoring of invader presence, cell traits and inheritance in single-cell linages. We found that direct interference, despite its dependence on various stochastic processes and poorly understood tug-of-war between replication of invading nucleic acids and degradation by CRISPR-Cas systems, is notably deterministic and efficient, with invader DNA clearance achieved in all cells within 1-3 generations. Conversely, the priming CRISPR-Cas response was highly variable, ranging from 2-30 generations before clearance. Our data show that direct interference and primed interference can in fact occur on comparable time scales, hence identifying the adaptation phase of priming as the origin of the variation.

Direct observations of the CRISPR-Cas action revealed several factors that impact CRISPR-Cas response variability. For direct interference we found that cells that cleared the target DNA earlier, grew and divided faster than the population mean. This might be explained by the fact that faster growth is known to reduce plasmid copy numbers ^52, 53^, while in slow growing cells plasmid maintenance mechanisms increase plasmid abundance ^54^. For priming the reverse was found. Cells that primed and cleared the target DNA earlier, grew more slowly, perhaps due to higher target DNA availability. The abundance of Cascade also plays a key role in direct interference and priming. We note that slower growing cells had higher Cascade abundance (Supplementary Fig. 17). Although single cell variability was not monitored, in line with our findings, spacer acquisition was found to occur more frequently during slower growth at late stationary phase ^37, 38^.

Our finding that target copy number influences the efficiency of spacer acquisition has implications for phage invasion. It suggests that one genome copy of a single virulent phage with an escape PAM may not lead to efficient CRISPR adaptation. However upon replication of the phage genome, it may become abundant enough, though at this point in time it is likely that primed interference with a new spacer cannot successfully eliminate a virulent phage before cell lysis ^27, 55, 56^. Despite this, it has been shown bioinformatically that priming by type I systems is widespread in nature ^57^, especially against temperate phages ^58^. Such events could occur due to low-level interference, in which a cell is able to simultaneously clear the invader while present as a single copy and acquire a spacer from the fragments produced. This would result in immunization of a single cell in the population, ultimately leading to a subpopulation of resistant cells that could limit further propagation of the same phage. Such a phenomenon may be more likely to occur when a defective phage infects the cell ^55^.

The variation existing between single cells in a population is remarkable. Stochasticity or noise in gene expression and cellular components has been demonstrated to play crucial roles in many cellular processes ^59–61^. We anticipate that the dynamics and heterogeneity of the CRISPR-Cas system, as studied here, play an important function in strategies that bacteria exploit and evolve in their continuous competition with phages, as well as with other species. For instance, CRISPR-Cas could contribute to bet-hedging strategies ^62^, in which subpopulations develop to combat changes in the environment, such as phage exposure. A distinct subpopulation in which Cascade is highly expressed could allow faster elimination of an invading phage, and subsequent re-population. This may in turn increase the fitness of the population, by reducing the overall burden of CRISPR-Cas expression and risk of autoimmunity ^30, 63^, and hence outcompete other bacterial strains. While such a strategy may not guarantee single cell survival, it is at large beneficial for the population as whole. Indeed, previous studies have shown CRISPR-Cas immunity in single cells acts to limit phage propagation throughout the population in an abortive infection like manner ^64–66^. On the other hand, the survival of only a subpopulation of cells may result in population bottlenecking and an overall loss of diversity ^67^. This may be disadvantageous in terms of spacer diversity, where it has been shown that populations containing a range of spacers are better able to combat and even facilitate the extinction of new invaders ^35, 47^.

While a number of studies have thoroughly investigated CRISPR-Cas systems through population and single molecule based experiments ^5, 7, 11, 30, 36–38, 64, 65, 68, 69^, these findings do not provide insight into the cell-to-cell variability. Our work has begun to bridge this gap demonstrating how important the dynamics of CRISPR-Cas systems are to their functioning and the outcome of populations facing a threat. Further investigation into different CRISPR-Cas types and classes, fluctuating environments ^70^, and conditions supporting the formation of subpopulations ^71^ will enhance the understanding of CRISPR-Cas dynamics on both the molecular and population scale.

## Methods

### Cloning

Plasmid pTU166 targeted by KD615 and KD635, was created by amplifying the streptomycin resistance cassette from pCDFDuet-1 with primers BN831 and BN832 to add a 5’CTT-PS8 tail. The backbone of pVenus was amplified using primers BN833 and BN834 and both products were restricted with KpnI and HindIII enzymes. Overnight ligation at 16 °C and transformation into DH5ɑ resulted in colonies selected to contain the plasmid. Plasmids pTU190 and pTU193 were created by PCR amplification of pTU166 using primer BN911 in combination with BN912 or BN891 respectively. Products were restricted with SalI, ligated and transformed into DH5a. Target plasmids pTU389 and pTU390 were PCR amplified from plasmid pTU265 a derivative of pVenus containing CFP using primers BN2278 in combination with BN2275 or BN2276 respectively. Products were restricted with NcoI, ligated and transformed into DH5a. All plasmids were confirmed by Sanger sequencing (Macrogen). All plasmids used are listed in Supplementary Table 1. Primers used are listed in Supplementary Table 2.

### Creation of strains KD615mCherry-Cas8e and KD635mCherry-Cas8e

Strains were created using lambda red homologous recombination ^72^. Plasmid pSC020, containing both Lambda red and the Cre-recombinase, was transformed by electroporation into strains KD615 and KD635. Strains were recovered at 30 °C for 1.5 h and plated on media containing 100 μg/ml ampicillin. Transformants were then grown overnight in liquid medium at 30 °C, with selection, and made competent the following day by inoculating 50 ml with 500 μl of overnight culture. Once the cells reached an OD600 of 0.2 a final concentration of 0.2% L-Arabinose (Sigma-Aldrich) was added and cells were grown for another 1.5 h and subsequently washed with pre-cooled 10% glycerol.

The mCherry-*cas*8*e* G-block (IDT) (Supplementary Table 3) based on the design used in ^3^ was resuspended with ddH20 to a concentration of 50 ng/μl and transformed into the competent cells by mixing 2 μl DNA with 50 μl of cells and recovering at 30 °C for 1.5 h. After recovery cells were plated undiluted with selection for kanamycin and ampicillin. PCR verified colonies were then grown in liquid culture with 1 mM IPTG at 37 °C to promote the loss of the kanamycin resistance cassette and pSC020. Individual colonies were screened for plasmid loss by patching each colony onto three plates containing no antibiotics, only kanamycin and only ampicillin. Colonies exhibiting no resistance were then PCR screened with primers (Supplementary Table 2) BN2204 and BN2205 for the presence of the mCherry-Cascade fusion. Strains were confirmed by Sanger sequencing (Macrogen).

### Growth conditions

All strain and plasmid combinations (Supplementary Table 1) used were grown at 37 °C, shaking at 180 rpm, prior to microscopy. To avoid autofluorescence under the microscope a minimal M9 media was used containing the following supplements; 2% glycerol (Sigma-Aldrich), 1X EZ Supplements (M2104 Teknova), 20 μg/ml uracil (Sigma-Aldrich), 1 mM MgSO4 (Sigma-Aldrich) and 0.1 mM CaCl2 (Sigma-Aldrich), from here on called M9 media.

### Microfluidic device

The device used was developed by D.J. Kiviet in the Ackermann lab and has been previously used in the Tans lab ^41^. The device contains a main flow channel 23.5 µm high and 200 µm wide that splits into two 100 µm wide flow channels of the same height. Perpendicular to these flow channels are wells with a height of 0.75 µm, widths of 1x 80 µm, 1x 60 µm, 2x 40 µm, 3x 20 µm, 3x 10 µm, 3x 5 µm and depths of 60 µm, 30 µm, 50 µm and 40 µm. These well sizes are repeated 5 times and are the location where the growth of microcolonies occurs during an experiment. The PDMS devices were made by casting them into an epoxy mould, a gift from D.J. Kiviet and the Ackermann lab.

The PDMS device was produced by mixing polymer and curing agent (Sylgard 184 elastomer, Dow Corning) in ratio of 1 mL of curing agent to 7.7 g of polymer. This mixture was poured into the epoxy mould and air bubbles were subsequently removed by use of a desiccator for 30 mins followed by baking at 80 °C for 1 h. After baking the device can be carefully removed from the mould with aid of a scalpel and holes were punched for liquid in-and outlets. For use under the microscope, the PDMS chip was covalently bound to a clean glass coverslip. This was done by treating both the PDMS and glass surface with 5-10 sweeps of a portable laboratory corona device (model BD-20ACV, Electro-Technic Products). After treatment the chip was placed carefully onto the glass slide and gently tapped to facilitate full contact between the PDMS and glass surface. Finally, the device was baked for another 1-2 h at 80°C and stored until the experiment was started.

### Loading and filling of microfluidic wells

Cells were initially grown overnight (for ∼ 12 h) at 37 °C, 180 rpm in 10 mL M9 media with antibiotic selection (streptomycin 50 µg/ml) for the target plasmid. The following day 500 µl of culture was passaged into fresh M9 medium (with selection for the target plasmid), approximately 3 h before microscope set up, and grown at 37 °C, 180 rpm. After 3 h of growth the cells were pelleted and resuspended in ∼ 30 µl.

To begin the experiment 2 µl of 0.01% Tween20 (dH2O) solution is slowly pipetted into the selected media lane to allow the removal of air and flow of liquid into the wells perpendicular to the media lane. Following this, 2 µl of concentrated bacterial culture was pipetted slowly into the same lane. Once liquid could be seen exiting at the opposite end of the media lane the syringes containing media (loaded on syringe pumps), the valve controller and the waste collection flasks were attached to the chip by metal connectors and polyethene tubing. Media was pumped into the chip at a flow rate of 0.5 mL/h allowing constant supply of nutrients to the cells. The rate of media flow was also important for removal of cells from the top of the well, to allow constant division and long-term tracking of cells located lower within the well.

### Media switches

All experiments were carried out with precise 37 °C temperature control and required the use of 2 different medias. For the first 12 h of the experiment (including loading of the chip) cells were grown in Media 1; M9 supplemented with both ahydrotetracycline (40 ng/ml) and Streptomycin (25 µg/ml) to induce the YFP and select for cells containing the target plasmid respectively. After 12 h of growth in the chip the media was switched via the valve controller (Hamilton, MPV valve positioner) to Media 2; M9 supplemented with anhydrotetracycline (40 ng/ml), 0.1% L-arabinose and 0.1 mM IPTG. This media change allowed removal of the selection for the target plasmid, continued induction of the YFP and induction of the CRISPR-Cas system after filling of the wells.

### Spacer acquisition detection from microfluidic chip output

Over the course of the experiment, the cells that flow out of the wells and subsequently the chip were collected in a sterile Erlenmeyer flask. The cells were then concentrated by centrifuging for 5 min at 2000 g. The supernatant was removed and cell were resuspended in 2 mL of M9 media. Colony PCR was performed 1 μl of culture using primers BN1530 and BN1531 (Supplementary Table 2) and the products were run on a 2% agarose gel at 100 V for 30 mins alongside the 100-1000 bp DNA Ladder (SmartLadder-SF, Eurogentec).

### Imaging and image analysis

For all time-lapse experiments, phase contrast images were acquired at 1 min intervals at a maximum of 2 positions. In experiments with a YFP target plasmid, fluorescent images were taken every 2 mins, with an exposure time of 500 ms. For experiments with a CFP target plasmid and the mCherry-Cascade fusion images were acquired every 4 mins with exposure times of 500 ms and 200 ms respectively. Images were acquired for the entire experiment including the first 12 hrs of growth. Cells were imaged with an inverted microscope (Nikon, TE2000), equipped with 100X oil immersion objective (Nikon, Plan Fluor NA 1.3), automated stage (Märzhäuser, SCAN IM 120 3 100), high power LED light source with liquid light guide (Sutter, Lambda HPX-L5), GFP, mCherry, CFP and YFP filter set (Chroma, 41017, 49008, 49001 and 49003), computer controlled shutters (Sutter, Lambda 10-3 with SmartShutter), cooled CMOS camera (Hamamatsu, Orca Flash4.0) and an incubation chamber (Solent) allowing temperature control. In order to obtain images with a pixel size of 0.041 µm an additional 1.5X lens was used. The microscope was controlled by MetaMorph software. A series of acquired phase contrast images were analyzed with a custom MATLAB (MathWorks) program, originally based on Schnitzcells software ^43^, adapted to allow for automated segmentation of cells growing in a well ^41^. Segmentation was inspected and corrected manually where necessary. All segmented cells were then tracked between frames using the pixel overlap between cells allowing the formation of lineage structures ^41^.

### Plasmid loss and clearance time detection using the fluorescent protein production rate

Before screening for plasmid loss, we detect cell death in lineages by applying a moving average filter to the cellular growth rate. If the cellular growth rate reached zero and did not recover again, the remainder of the fluorescence time series after this point was excluded from the analysis. For each lineage, we computed the fluorescence production rate of the plasmid-encoded fluorophore from a cell’s total fluorescence, cell area, cellular growth rate, and the rate of photobleaching of the fluorophore ^44^. As there is always some amount of residual fluorescence produced by the cells, we selected an appropriate threshold for plasmid loss detection from the upper values of the distribution of production rates of plasmid-free cells. To detect plasmid loss in individual lineages we applied a moving average filter to the fluorescence production rate and detected the first instance of the production rate reaching a value below the threshold. This plasmid loss time (PLT) can be seen as an upper bound estimate, as some processes (transcription, translation, fluorophore maturation) still carry on for some time after the last plasmid has been cleared but could not be measured in our set up. The onset of the clearance time (CT), which signifies the start of the destruction of all plasmids through interference and ends at the plasmid loss time (PLT), is difficult to detect in individual lineages due to the naturally occurring fluctuations in the fluorescence production rate. To determine this quantity, we align all plasmid loss lineages at the PLT and compute the average trend. The CT per experimental condition is approximated as the duration from the point where the average production rate starts to decrease until the PLT.

### Sister and cousin statistics

For each lineage that lost the plasmid, we wanted to compare the probability of loss in an unrelated cell and in a related cell. For related cells we counted the frequency of loss and non-loss in sister and cousin cells of the loss cell, but only if the sister or cousin divided (contained a complete cell cycle). For unrelated cells we counted the total number of loss events (*i*) that occurred throughout the cell cycle of the related cell. For each loss event we counted how many cells (*c*_*i*_) still contained the plasmid up to that point. The probability of plasmid loss happening in an unrelated cell during the lifecycle of the related cell was subsequently calculated recursively using the following equations:

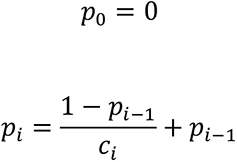

Where *p*_*i*_ is the probability of loss occurring within an unrelated cell given *i* plasmid loss events occurred within the cell cycle of the related cell and *c*_*i*_ stands for the number of cells still containing the plasmid at the same time as the *i*-th plasmid loss event.

### Cascade copy number determination

The control strain KD614 mCherry-Cas8e containing plasmid pTU265 (Supplementary Table was prepared and loaded into the microfluidic chip as above. After 12 h a sterile tube was connected to the waste tubing and output from the chip was collected for 30 mins. The media was then switched to induce Cascade. Approximately 5 h after induction when Cascade levels are considered to be stabilized the output from the chip was again collected for 30 mins. To improve counting, cells were subsequently fixed with 2.5% paraformaldehyde solution at 22 °C for 45 mins ^73^. Slides were cleaned by sonication in subsequent steps with MilliQ, acetone and KOH (1M). Next, 1 % agarose pads containing the M9 medium were prepared and hardened between two slides within 20 mins of measuring to prevent desiccation. The fixed cells were then spun down and resuspended in 5 µl of which 1 µl was pipetted onto a pre-prepared agarose pad.

The cells were imaged using a TIRF microscope (Olympus IX81; Andor Ixon X3 DU897 EM-CCD camera) using a high power 561 nm laser, which quickly bleached most mCherry molecules within a couple of frames. Intensity of single molecules were measured with Thunderstorm starting from the thirtieth frame ^74^. The total cell fluorescence was measured by segmenting the cells from the phase contrast image and sum fluorescence counts of all cell pixels (with background subtracted). The copy number was calculated by dividing the total cell fluorescence in the first frame by the average fluorescence intensity of the single molecules. We could then calculate the Cascade concentration ∼200 Cascade molecules/µm^2^ by dividing the population average of the mean summed RFP per cell by this copy number, which was applied to the cells in our time-lapse data.

### Model implementation

Stochastic simulations were performed using the adapted *Extrande* algorithm ^75^ implemented in C++. Each data point in Fig. 5i-j and Fig. 6a-c was obtained from 100 simulated experiments of up to 10^4^ min. The population size of each simulation was fixed at 100 cells. See Supplementary Methods for model details and parameters.

## Acknowledgements

The authors would like to thank Martijn Wehrens for his help and advice throughout the project and all of the members of the Tans and the Brouns groups for input during group discussions. We acknowledge the group of Konstantin Severinov for the gift of strains KD615 and KD635. R.E.M. is supported by the Frontiers of Nanoscience (NanoFront) program from NWO/Ministry of Education (OCW). C.F. received funding from FET-Open research and innovation actions grant under the European Union’s Horizon 2020 research and innovation programme (CyGenTiG; grant agreement 801041). S.J.J.B. has received funding from the European Research Council (ERC) CoG under the European Union’s Horizon 2020 research and innovation programme (grant agreement No. [101003229]) and from the Netherlands Organisation for Scientific Research (NWO VICI; VI.C.182.027).

## Author contributions

R.E.M., S.J.J.B. and S.J.T. and conceived the project, R.E.M., J.V., J.V.L. and V.K. performed the experiments. E.M.K., J.V., R.E.M. and F.B. analysed the data. E.M.K. and C.F. performed the modelling, R.E.M., E.M.K., S.J.J.B., S.J.T., and C.F., wrote the manuscript with input from all authors.

## Competing interests

The authors declare no competing interests.

## Reporting summary

Further information on research design is available in the Nature Research Reporting Summary linked to this article.

## Data availability statement

The data that support the findings of this study are available from the corresponding authors on request.

## Code availability statement

Data analysis was performed using custom MATLAB scripts, which can be found at https://github.com/TansLab/Tans_Schnitzcells. Scripts for lineage analysis and plotting were implemented in MATLAB and are available upon request. An implementation of the agent-based model in C++ is available at https://git.wur.nl/Biometris/articles/CRISPR_ABM.

## Additional information

## Notes

### Competing Interest Statement

The authors have declared no competing interest.

