## Supplementary Information for "Single Cell Variability of CRISPR-Cas Interference and Adaptation"

**Supplementary Figures**

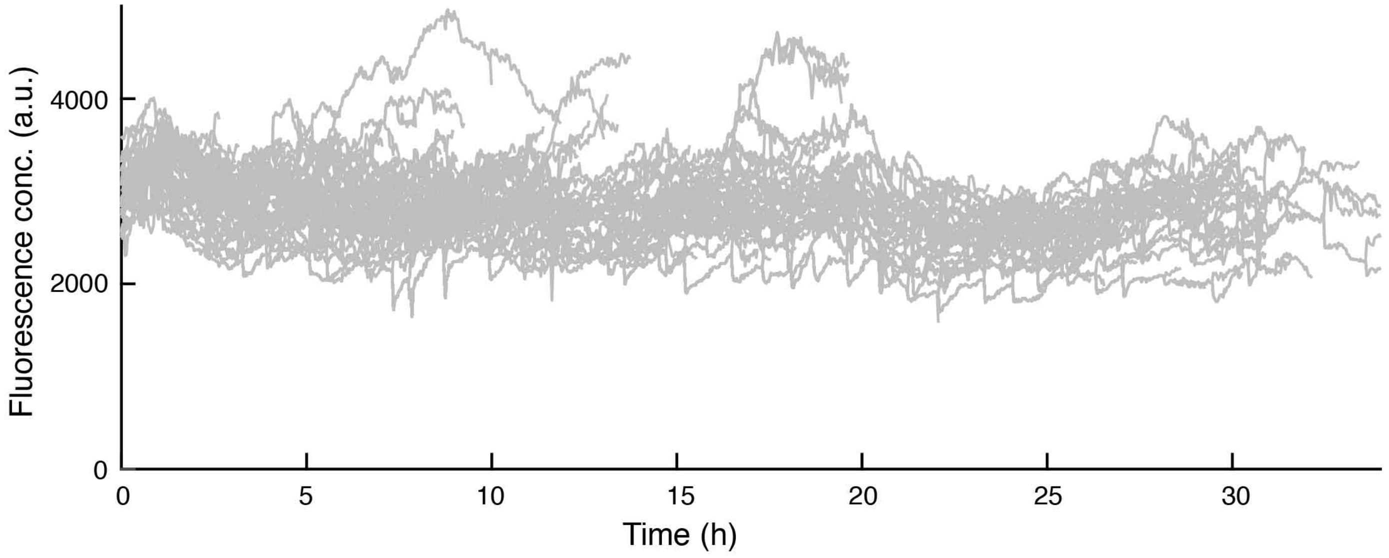


**Supplementary Fig. 1| Plasmid loss is CRISPR-dependent**

The YFP fluorescence traces in arbitrary units (a.u.) of the WT strain harboring pControl a plasmid with no target for the CRISPR-Cas system. Time-lapse imaging was carried out for 35 hours post induction of the *cas* genes, and showed no plasmid loss.

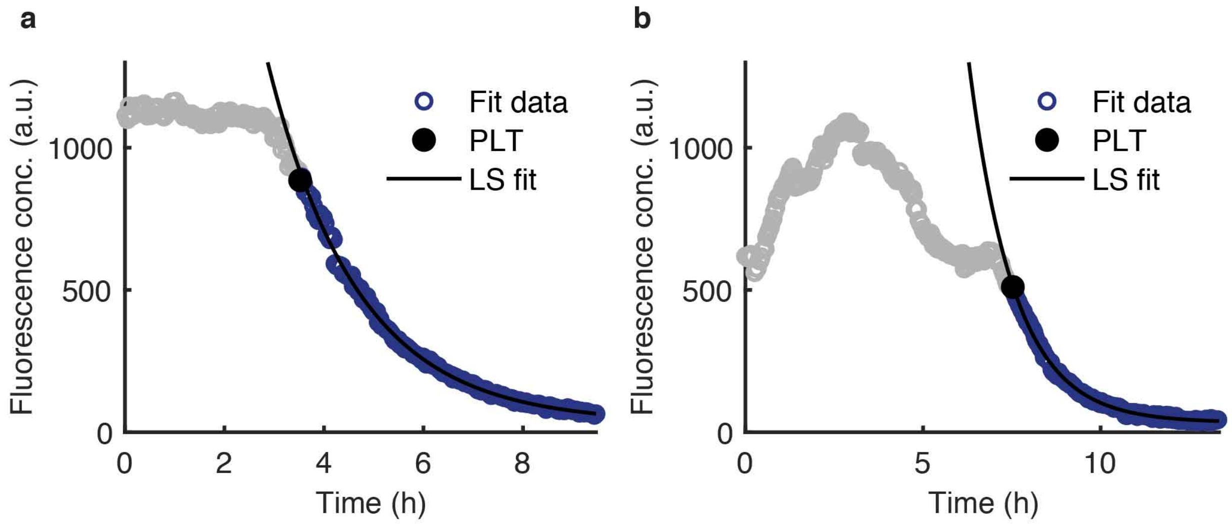


**Supplementary Fig. 2| Decay of YFP fluorescence in both direct interference and priming follows exponential decay**

The fluorescence concentration (open circles) of (**a**) direct interference and (**b**) priming lineages can be described by exponential decay. This was evaluated by performing a least-squares (LS) fit of the fluorescence concentration data (purple open circles) after the moment of plasmid loss (PLT, black circle) to an exponential curve (LS fit, black line).

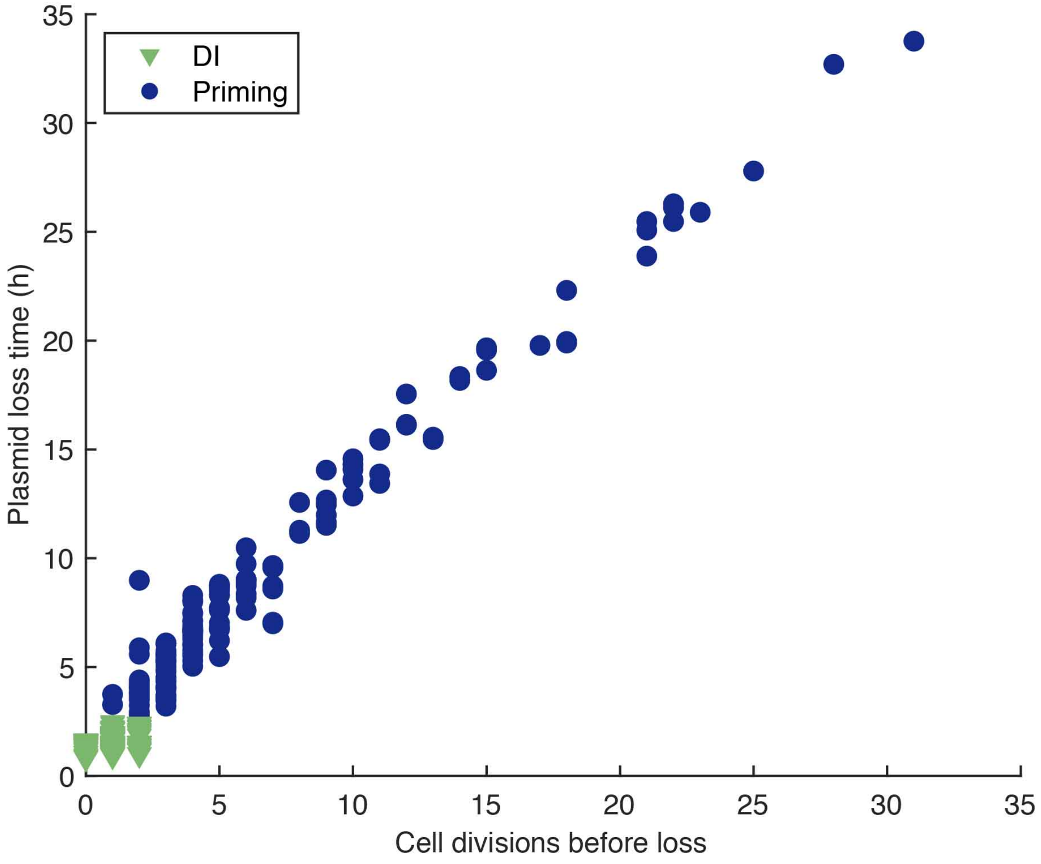


**Supplementary Fig. 3| Cell divisions before plasmid loss highly correlates with plasmid loss time**

The number of cell divisions from the moment of induction until plasmid loss are plotted against the PLT in hours. Both consensus target clearance by direct interference (green) and mutant target clearance by priming (blue) are shown.

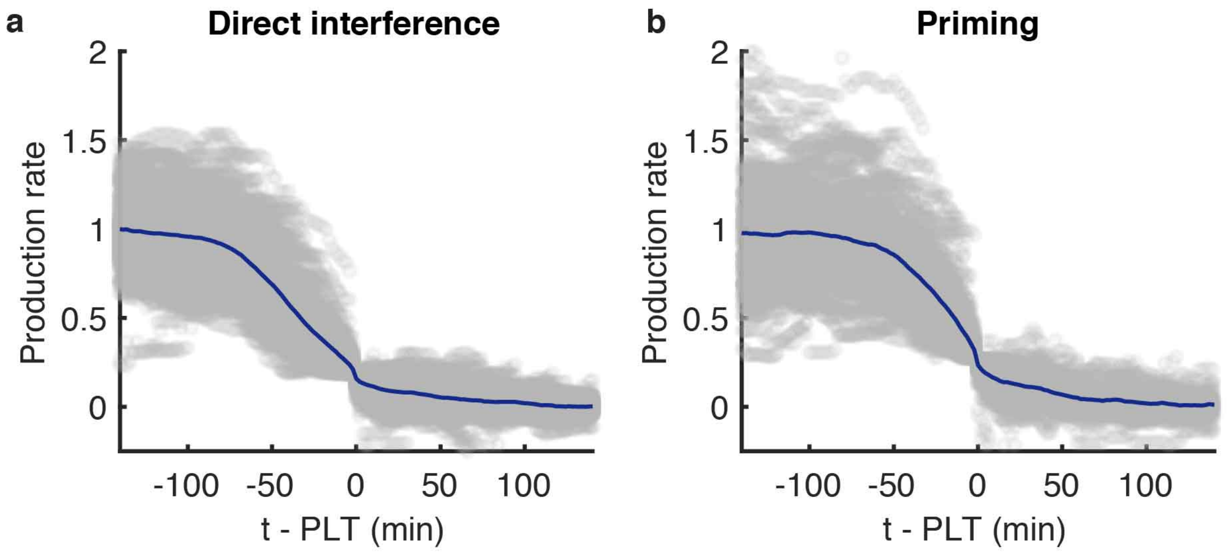


**Supplementary Fig. 4| Plasmid loss during Direct interference and primed interference processes occur on a comparable timescale**

All production rate traces (grey) starting from 140 minutes prior to the detected plasmid loss time PLT from (**a**) direct interference and (**b**) priming were aligned at the PLT (t-PLT=0) and the average trend (navy) normalized for comparison. From the average trend, we estimate the clearance time (CT), time taken from the initiation of plasmid clearance until elimination of all copies, to be in the order of 60 minutes for both direct interference and priming from the onset of the production rate decrease.

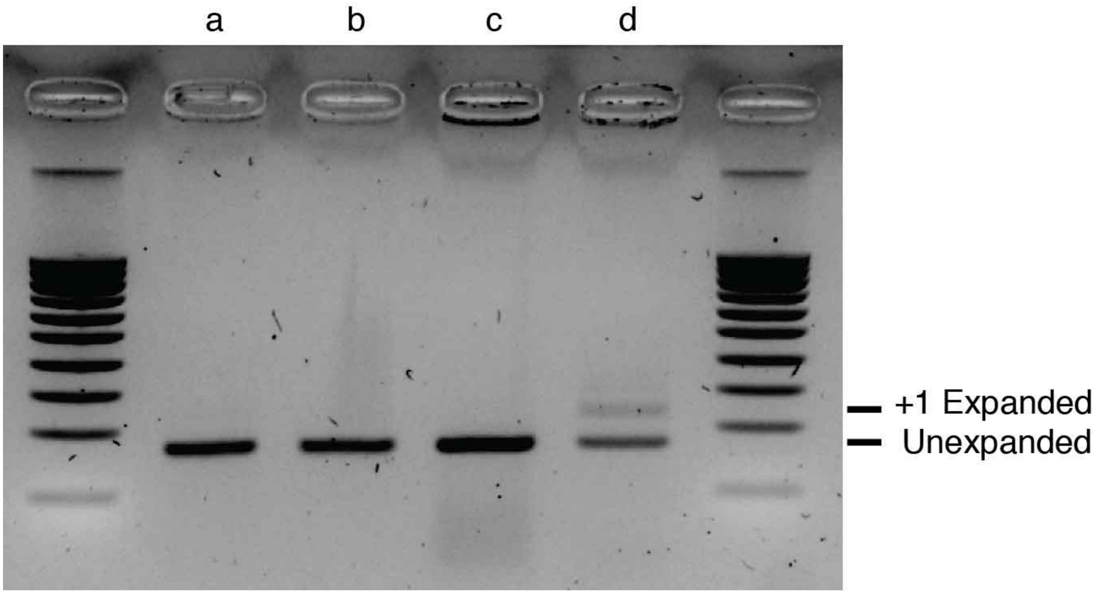


**Supplementary Fig. 5| Spacer acquisition was only seen in the WT strain in the presence of a mutated PAM triggering priming**

Cells from the chip output were collected in a flask for each experiment and the CRISPR arrays were screened for expansion due to spacer acquisition by PCR amplification using primers BN1530 + BN1531 (Supplementary Table 2). The gel shows PCR amplified CRISPR arrays from each experiment **a,** WT + pControl (Control) **b,** Δ*cas*1,2 + pTarget (direct interference) **c,** Δ*cas*1,2 + pMutant **d,** WT + pMutant (priming). The presence of a larger band indicates array expansion and therefore successful spacer acquisition.

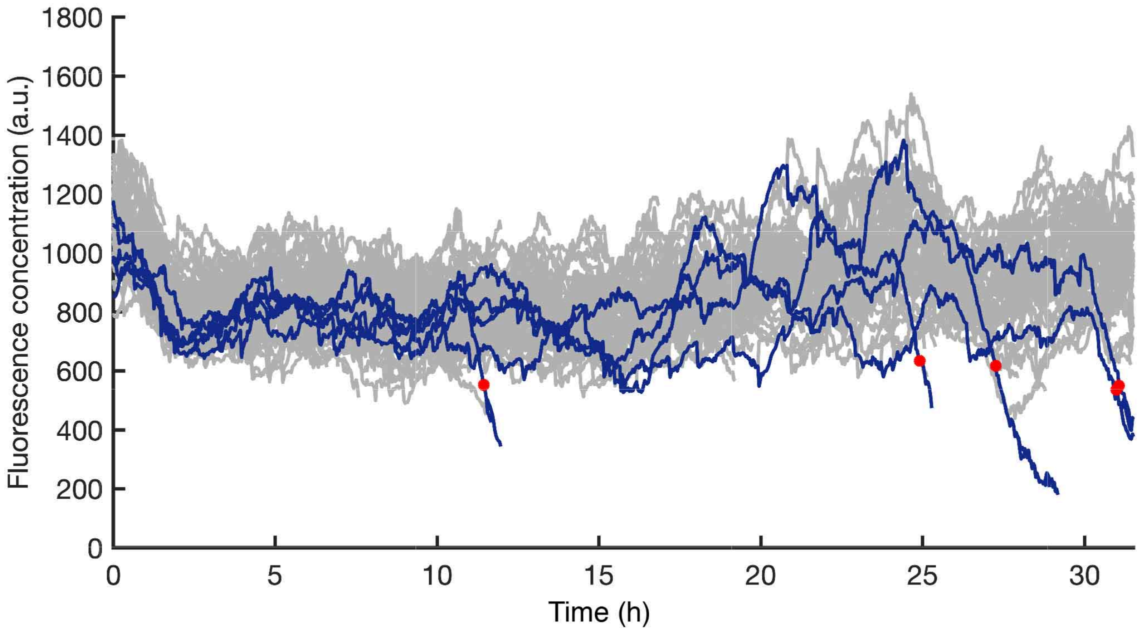


**Supplementary Fig. 6| In the absence of Cas1 and Cas2 clearance of a target with a non-consensus PAM mutant occurs rarely**

The YFP fluorescence of the $\Delta$*cas*1,2 strain containing pMutant was imaged for 34 hours after induction. Lineages that were able to clear the plasmid are highlighted in blue with the red dot indicating the moment of detection. 1.4% of lineages (5 unique events, red dot) cleared the plasmid.

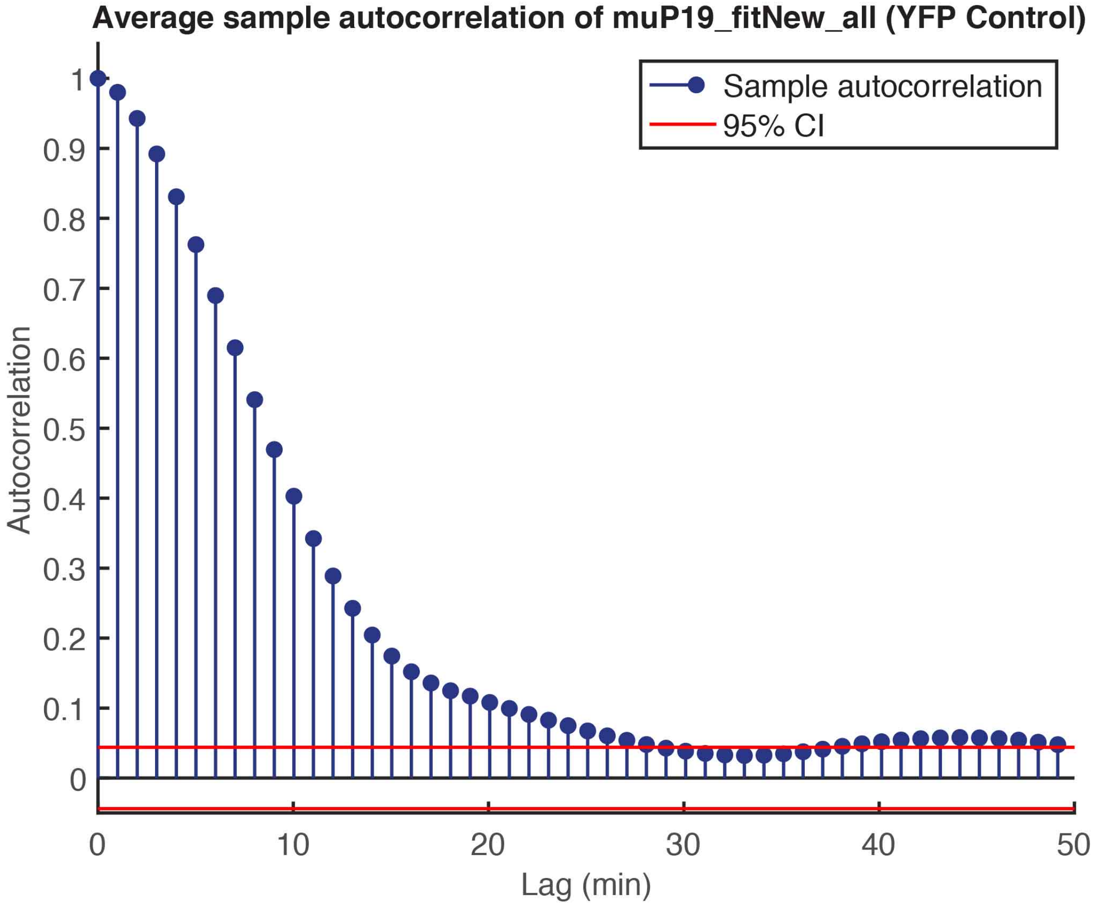


**Supplementary Fig. 7| Autocorrelation time of cellular growth rate**

The autocorrelation time was calculated for the cellular growth rate of the WT strain containing pControl by averaging the autocorrelation of cell growth as a function of time in individual lineages. After 10 mins the autocorrelation of cellular growth has decreased to 0.4. After approximately 30 minutes the autocorrelation has decreased to zero, as indicated by 95% confidence intervals (red lines).

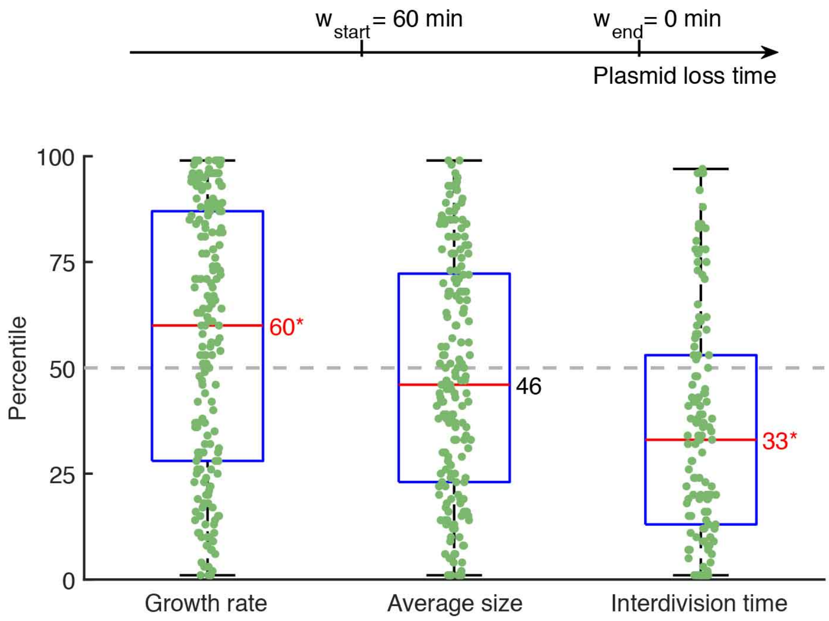


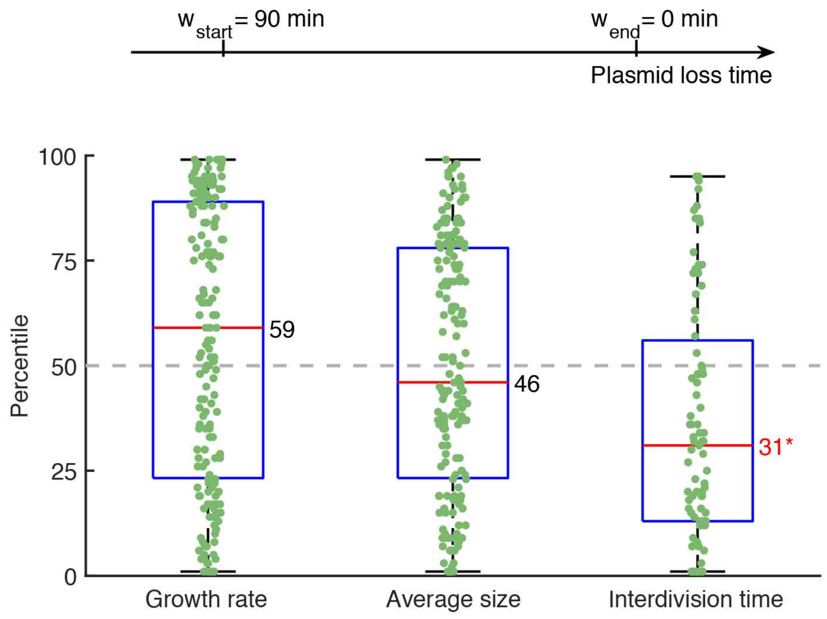


**Supplementary Fig. 8| Growth rate, cell size and interdivision time of direct interference with different lookback windows**

Boxplots of growth rate, average cell size and interdivision time presented as the percentile rankings of all plasmid loss lineages (green) that cleared a known target via direct interference. The cell feature of interest (e.g. growth rate) was averaged over a lookback window chosen in relation to the time from plasmid loss of the lineage of interest. The same cell feature was then averaged for all non-loss lineages in the population at that same moment. The cell feature of interest was then ranked amongst the non-loss population as a percentile. We considered lookback windows of 60 minutes prior to plasmid loss (top) and

90 minutes prior to plasmid loss (bottom). The median percentile ranking of loss lineages is indicated by a red line and black text, categories in which this value was significantly different from a ranking in the $50^{th}$ percentile ($p$-value<0.05) are indicated in red text followed by an asterisk.

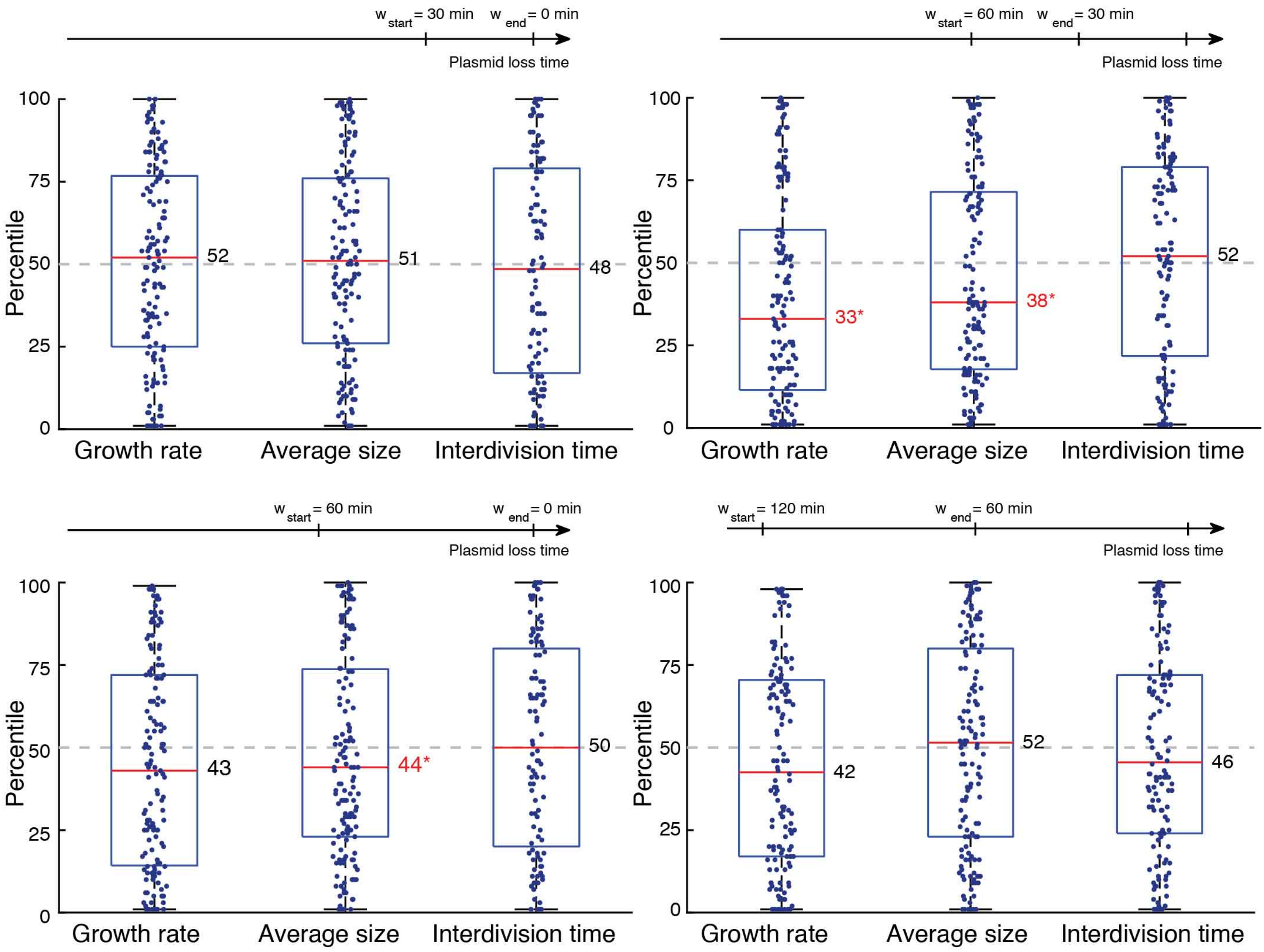


**Supplementary Fig. 9| Growth rate, cell size and interdivision time of priming with different lookback windows**

Boxplots of growth rate, average cell size and interdivision time presented as the percentile rankings of all plasmid loss lineages (navy) that cleared a known target via priming. The cell feature of interest (e.g. growth rate) was averaged over a lookback window chosen in relation to the time from plasmid loss of the lineage of interest. The same cell feature was then averaged for all non-loss lineages in the population at that same moment. The cell feature of interest was then ranked amongst the non-loss population as a percentile. We considered a range of lookback windows. The median percentile ranking of loss lineages is indicated by a red line and black text, categories in which this value was significantly different from a ranking in the 50$^{th}$ percentile (p-value <0.05) are indicated in red text followed by an asterisk.

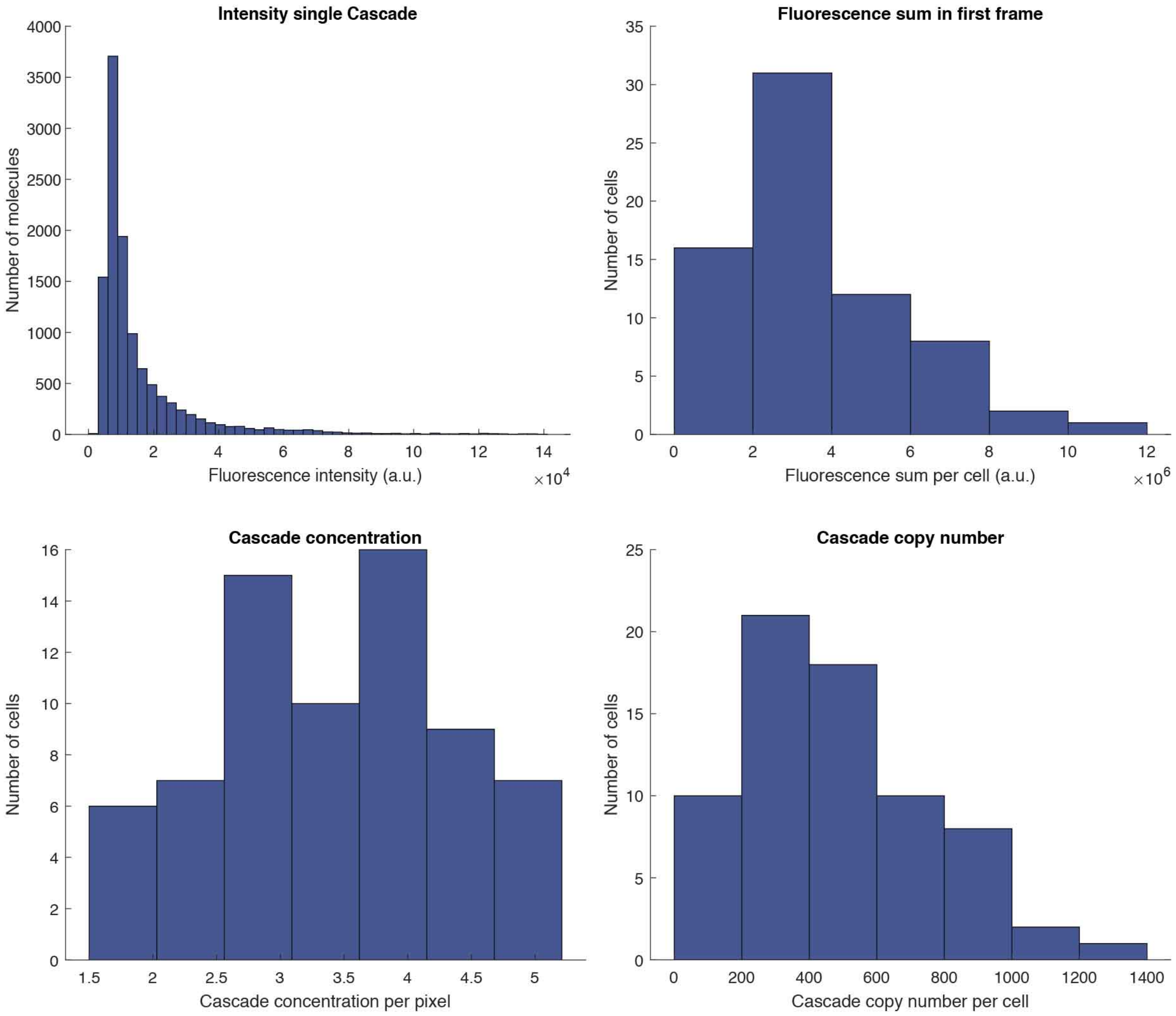


**Supplementary Fig. 10| Cascade copy number determination**

**a,** The fluorescence sum (RFP) of each cell in the first frame was determined. **b,** The RFP molecules were then bleached until it was possible to determine the fluorescence intensity of a single molecule (representing a single Cascade). **c,** The Cascade copy number per cell was then determined by dividing the average fluorescence sum by the average intensity of a single Cascade molecule. **d,** The Cascade concentration per pixel was determined by dividing the fluorescence sum by the area of the cell in pixels.

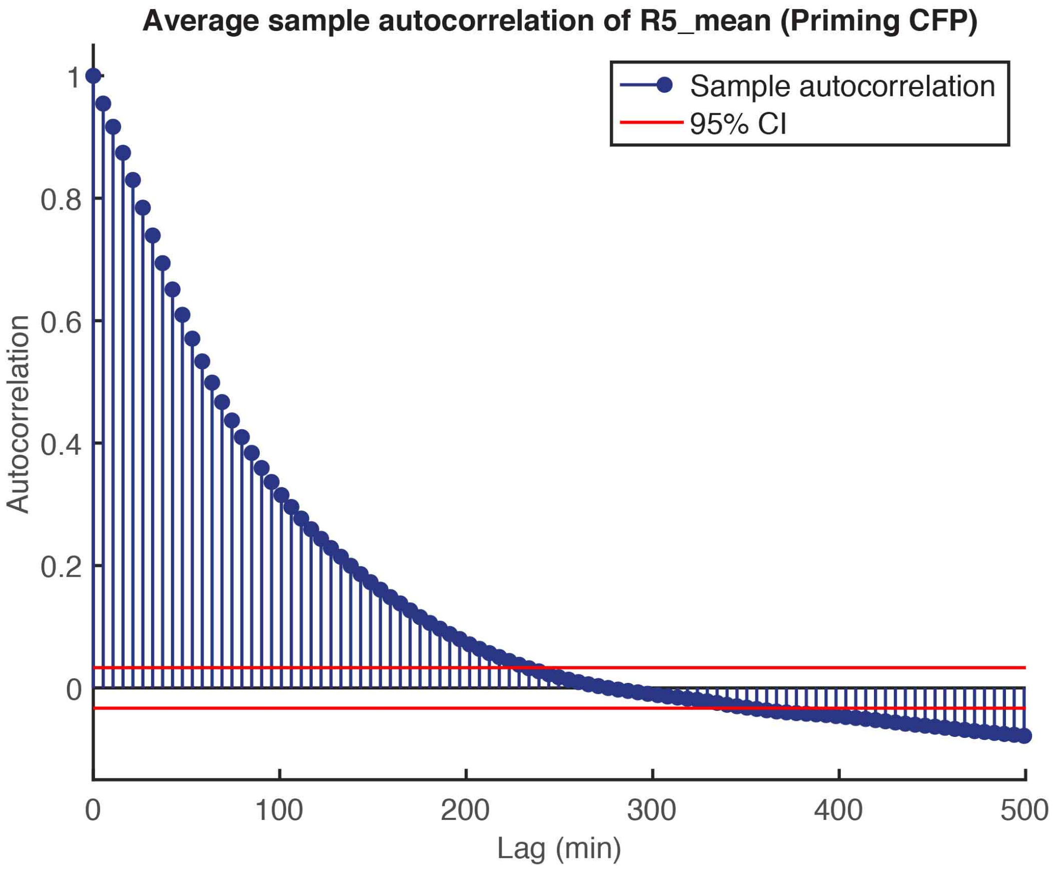


**Supplementary Fig. 11| Autocorrelation of RFP (Cascade) concentration**

The autocorrelation was calculated by averaging over the autocorrelation of RFP concentration of the WT-mCherry strain in individual lineages. After approximately 200 minutes the autocorrelation has decreased to zero, as indicated by 95% confidence intervals (red lines). The long decay time of the autocorrelation function indicates that Cascade protein levels fluctuate on a time scale longer than the cell cycle.

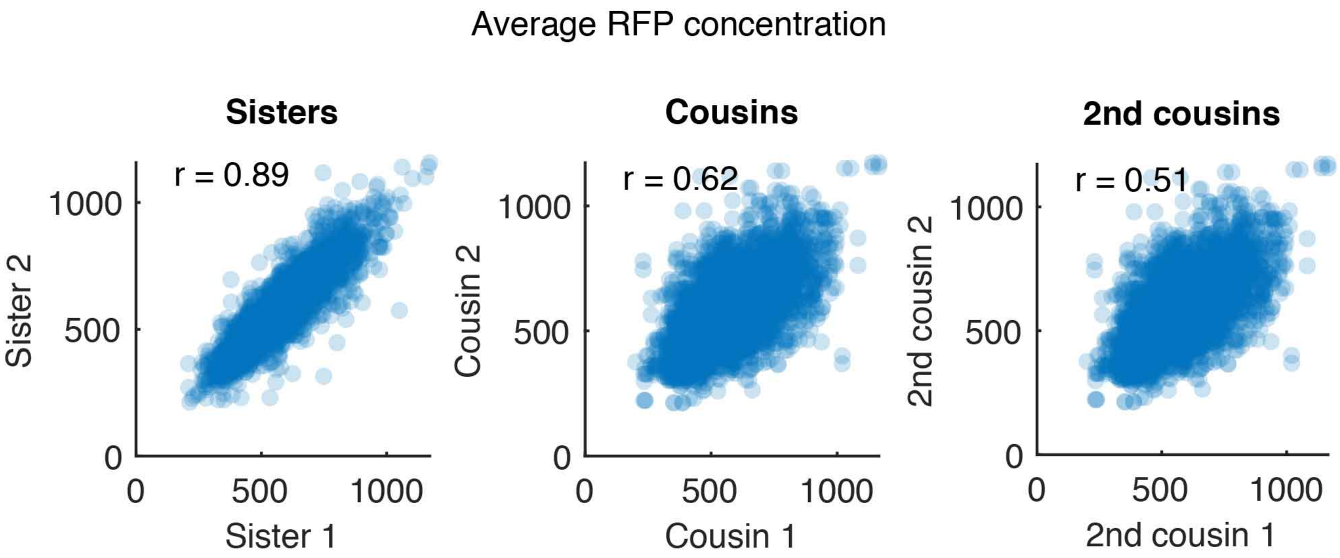


**Supplementary Fig. 12| Correlation of RFP levels between cells related as sisters, cousins, and second cousins**

The levels of RFP (Cascade) are strongly correlated between sister, cousins, and second cousins. The correlation coefficient r decreases as the cells become less closely related.

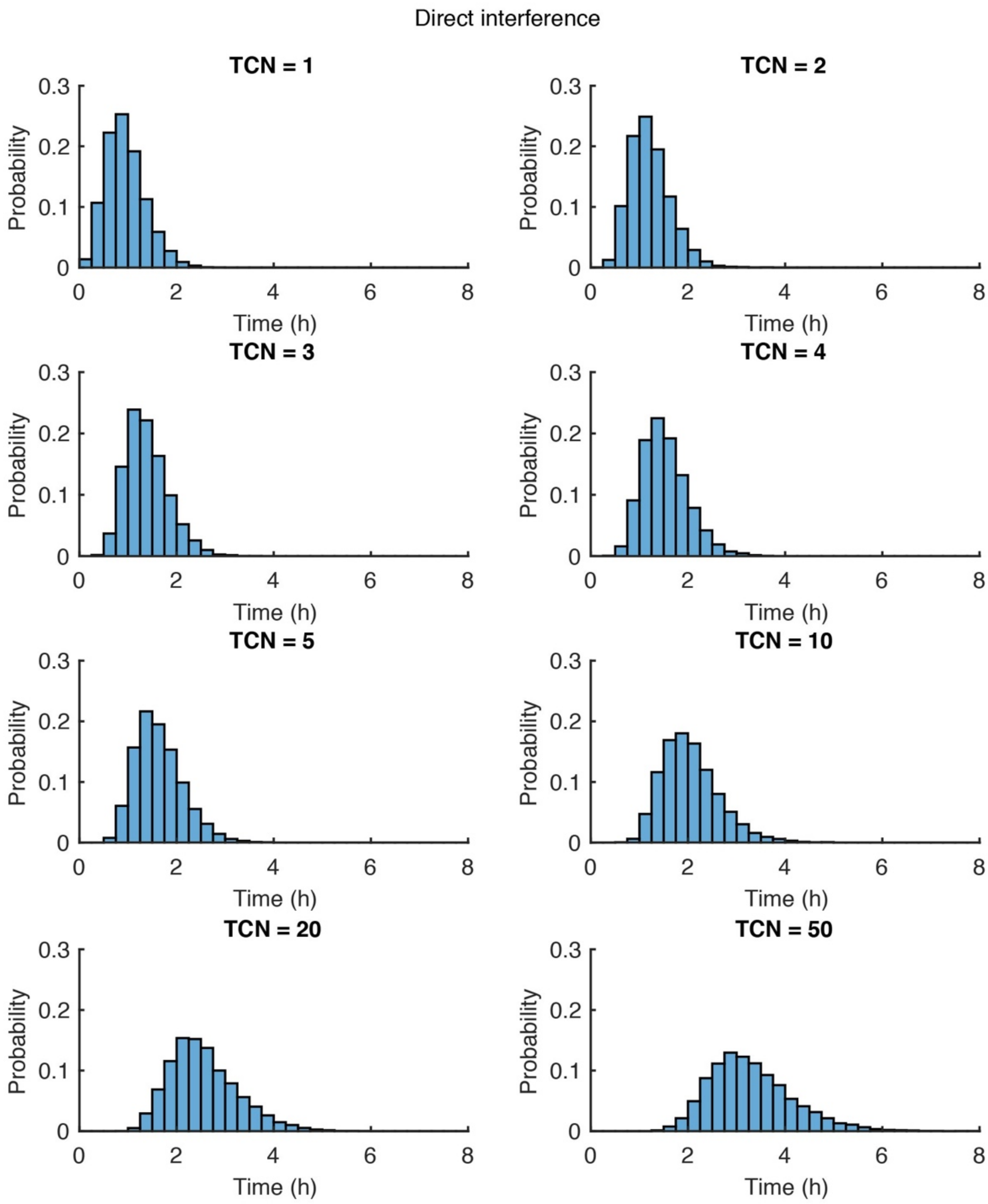


**Supplementary Fig. 13| Distribution of target loss times resulting from simulations of the direct interference condition for average target copy numbers (TCN) per cell ranging from 1-50**

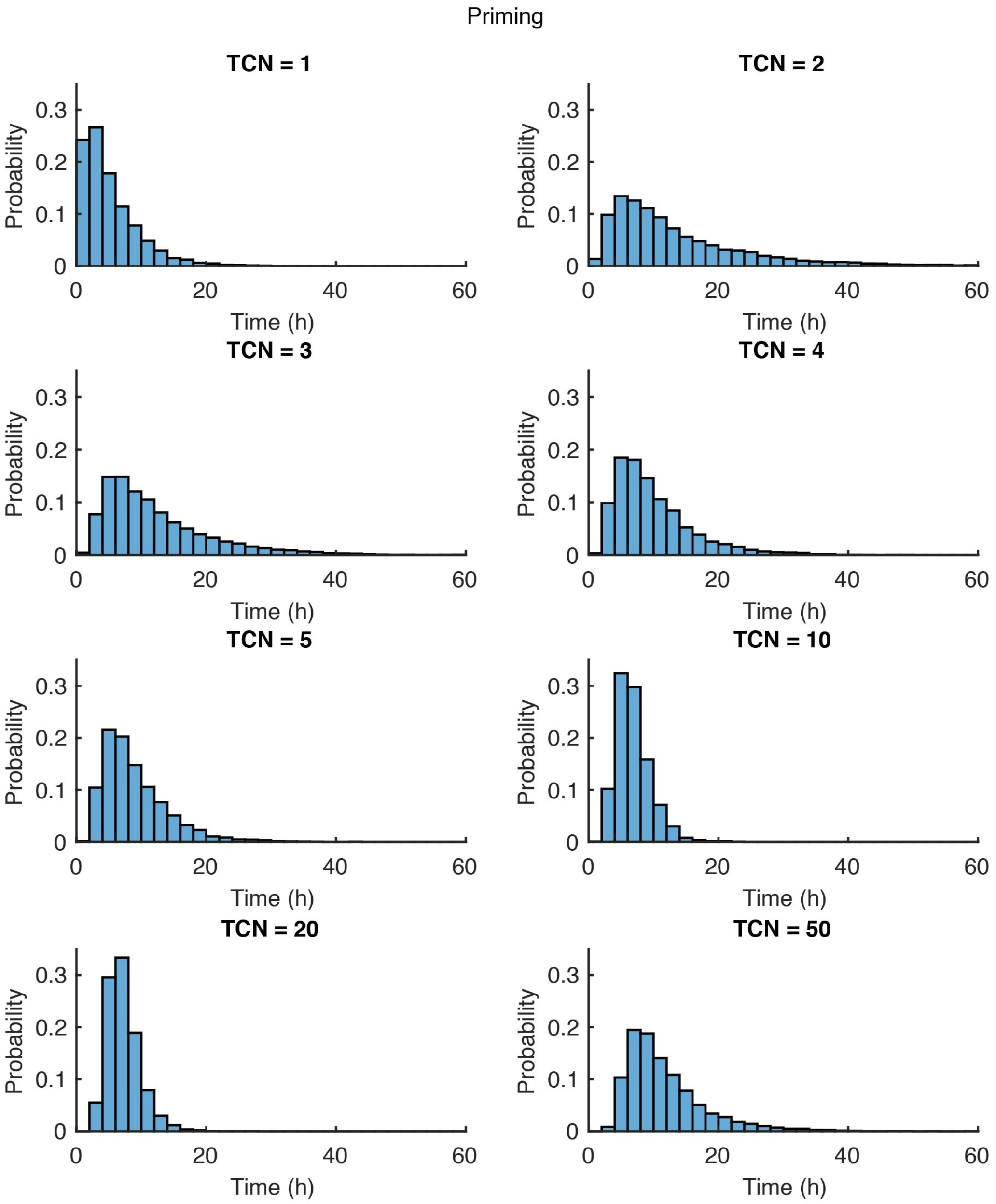


**Supplementary Fig. 14| Distribution of target loss times resulting from simulations of the priming condition for average target copy numbers (TCN) per cell ranging from 1-50**

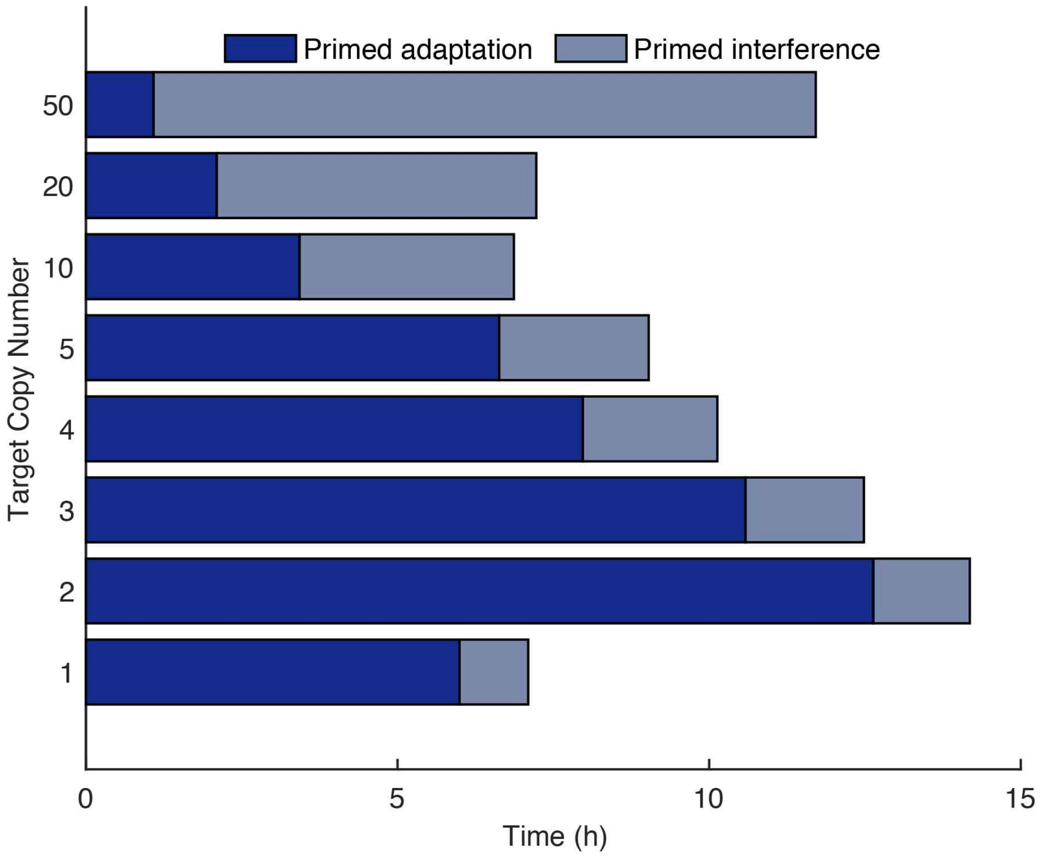


**Supplementary Fig. 15| Target loss time as a function of the target copy number (TCN) as computed from simulated trajectories by the agent-based model for the priming condition**
Bar charts representing the time spent on primed adaptation (navy) and primed interference (grey) for cells clearing targets through priming with an average plasmid copy number ranging from 1-50.

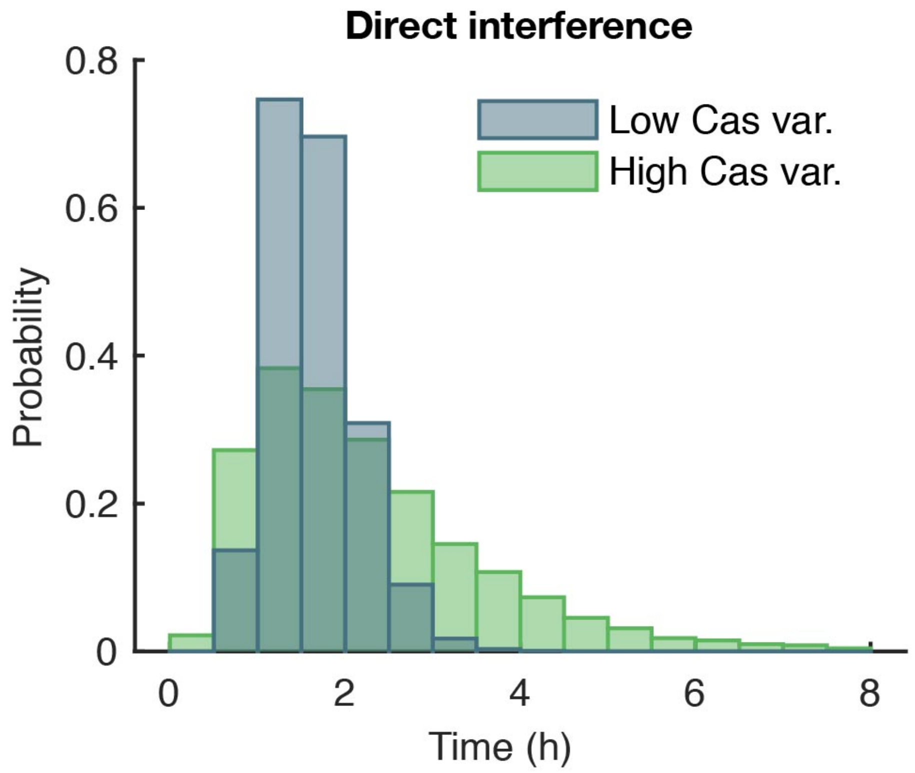


**Supplementary Fig. 16| Distribution of plasmid loss times in direct interference for high and low variability in Cascade concentration**

Target loss time distribution for two different levels of Cascade concentration variability resulting from simulated trajectories of the direct interference condition. At low variability (blue) Cascade proteins are produced in frequent, small bursts, whereas at high variability (green) proteins are synthesized more sporadically in large bursts (100-fold increase), keeping average Cascade concentration constant. The variability of PLT interference times for high Cascade variability increases as compared to low Cascade variability.

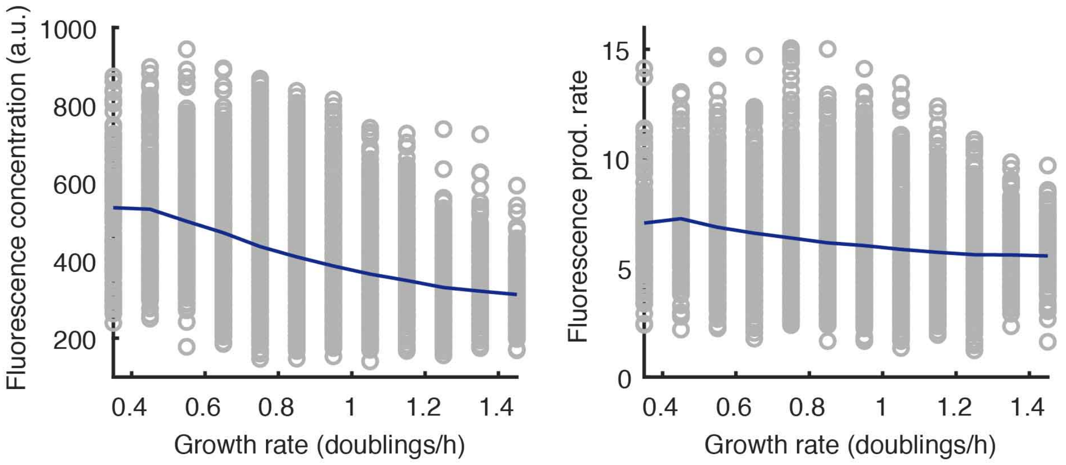


**Supplementary Fig. 17| Slower growing cells have higher RFP (Cascade) concentrations**

Cascade concentration (left) and Cascade production rate (right) show an inverse relationship with cellular growth rate (grey circles), revealing slower growing cell on average (navy line) have a higher concentration of Cascade.

**Supplementary Table 1. Strains and plasmids used in this study**

| **Strain** | Description | Source |
|---|---|---|
| KD615 | *E. coli* K12, F+, araBp8-*cse*1, lacUV-*cas*3, CRISPR I R-SP8-R, ΔCRISPR II+III | (Musharova et al., 2019) |
| KD635 | *E. coli* K12, F+, araBp8-*cse1,* lacUV-*cas*3, CRISPR I R-SP8-R, Δ*cas*1,2, ΔCRISPR II+III | (Musharova et al., 2019) |
| KD615mCherry-Cas8e | *E. coli* K12, F+, araBp8-*cse1,* lacUV-*cas*3, CRISPR I R-SP8-R, ΔCRISPR II+III, mCherry-*cas*8*e* | This study |
| KD634mCherry-Cas8e | *E. coli* K12, F+, araBp8-*cse1,* lacUV-*cas3,* CRISPR I R-SP8-R, Δ*cas*1,2, ΔCRSIPR II+III, mCherry-*cas*8*e* | This study |
| **Plasmid** | | |
| pTarget (pTU166) | pSC101, StrepR, TetR mVenus PS8 flanked by 'CTT' PAM | This study |
| pMutant (pTU190) | pSC101, StrepR, TetR mVenus PS8 flanked by 'CGT' PAM | This study |
| pControl (pTU193) | pSC101 ori, StrepR, TetR-mVenus, no target | This study |
| pVenus | pSC101 ori, KanR, mVenus-YFP | Bokinsky lab |
| pCDFDuet-1 | pCloDF13 ori, StrepR | Lab collection |
| pTU265 | pSC101, StrepR, TetR-Cerulean, no target | This study |
| pTU389 | pSC101, StrepR, TetR-Cerulean, PS8 flanked by 'CGT' PAM | This study |
| pTU390 | pSC101, StrepR, TetR-Cerulean, PS8 flanked by 'CTT' PAM | This study |
| pSC020 | Derivative of pKD46 containing Lamda red and the Cre-recombinase | This study |

**Supplementary Table 2. Oligonucleotides used in this study**

| Name | Description | Sequence |
|---|---|---|
| BN831 | Streptomycin resistance and PS8 insertion into pVenus, Fw | TTTTGGTACCTTATTTGCCGACTACCTTGGTGATCTC |
| BN832 | Streptomycin resistance and PS8 insertion into pVenus, Rv | TTTTAAGCTTAAAAGTGCCACTTGCGGAGACCCGGTCGTCAG**CTT**ACATTCAAATATGTATCCGCTC |
| BN833 | Backbone amplification pVenus, Rv | TTTTGGTACCGGACTCTGGGGTTCGAG |
| BN834 | Backbone amplification pVenus, Fw | TTTTAAGCTTCGAAACGATCCTCATCCTG |
| BN891 | Streptomycin resistance insertion (no target), Rv | TTTTAAGCTTACATTCAAATATGTATCCGCTC |
| BN911 | Modify pTU166 PAM universal, Rv | TTTTGTCGACACATTCAAATATGTATCCGCTCATGAGAC |
| BN912 | Modify pTU166 CTT PAM to CGT | TTTTGTCGAC**ACG**CTGACGACCGGGTC |
| BN1494 | To amplify pTU193 Backbone minus yfp, Rv | TTTCTCGAGTAAGGATCTCCAGGCATC |
| BN1495 | To amplify pTU193 Backbone minus yfp, Fw | TTTCTCGAGTAAGGATCTCCAGGCATC |
| BN1507 | To amplify Cerulean from p15A, Fw | TTTGAATTCCAGAATTCAAAAGATCTAGGAGG |
| BN1508 | To amplify Cerulean from p15A, Rv | TTTCTCGAGAGGATCCTTATTTATACAGCTCATCC |
| BN1513 | To check Cerulean insertion and confirm pTU265 by sequence, Fw | CCTCATTAAGCAGCTCTAATGCGCTG |
| BN1530 | To screen for CRISPR array amplification, Fw | GGTTTGAAAATGGGAGCTCG |
| BN1531 | To screen for CRISPR array amplification, Rv | GTTACATTAAGGTTGGTGGGTTG |
| BN2202 | To amplify mCherry-Cas8e gblock, Fw | ACAGAATCTGGATGGATGG |
| BN2203 | To amplify mCherry-Cas8e gblock, Rv | CTGATCTCTACTGCAGTATAGC |
| BN2204 | Screen for mCherry-*cas8e* knock in, Fw | GCGCTTGCACTTAATCGC |
| BN2205 | Screen for mCherry-*cas8e* knock in, Rv | ACCAGCAGTGCTAAAGCG |
| BN2206 | Screen for mCherry-*cas8e* knock in, Fw | CTTTCCGTCCGGTGTCAGG |
| BN2275 | Insertion PS8 CGT PAM into pTU265, Fw | TTTCCATGGAAAAGTGCCACTTGCGGAGACCCGGTCGTCAG**CGT**ACATTCAAATATGTATCCGCTCAT |

| | | |
|---|---|---|
| BN2276 | Insertion PS8 CTT PAM into pTU265, Fw | TTTCCATGGAAAAGTGCCACTTGCGGAGACCCGGTCGTCAG**CTT**ACATTCAAATATGTATCCGCTCAT |
| BN2278 | Insertion of PS8 and PAM universal, Rv | TTTCCATGGCCTCATCCTGTCTCTTGATC |

***PAM sequences are indicated in bold, and restriction sites are underlined.**

**Supplementary Table 3. Synthetic DNA G-block used in this study**

| Name | Sequence |
|---|---|
| *cas8e*-mCherry insert | ACAGAATCTGGATGGATGGGTCTGGCAGGGTAACAGTATTGTTATTACCTATACA<br>GGGGATGAAGGGATGACCAGAGTCATCCCTGCAAATCCCAAATAACCTGGAGCT<br>GCAGATACCGTTCGTATAATGTATGCTATACGAAGTTATAGATCTCTATTTGTTTAT<br>TTTTCTAAATACATTCAAATATGTATCCGCTCATGAGACAATAACCCTGATAAATGC<br>TTCAATAATATTGAAAAAGGAAGAGTATGAGCCATATTCAACGGGAAACGTCTTG<br>CTCTAGGCCGCGATTAAATTCCAACATGGATGCTGATTTATATGGGTATAAATGGG<br>CTCGCGATAATGTCGGGCAATCAGGTGCGACAATCTATCGATTGTATGGGAAGCC<br>CGATGCGCCAGAGTTGTTTCTGAAACATGGCAAAGGTAGCGTTGCCAATGATGTT<br>ACAGATGAGATGGTCAGACTAAACTGGCTGACGGAATTTATGCCTCTTCCGACCAT<br>CAAGCATTTTATCCGTACTCCTGATGACGCATGGTTACTCACCACTGCGATCCCCG<br>GGAAAACAGCATTCCAGGTATTAGAAGAATATCCTGATTCAGGTGAAAATATTGT<br>TGATGCGCTGGCAGTGTTCCTGCGCCGGTTGCATTCGATTCCTGTTTGTAATTGTC<br>CTTTTAACAGCGACCGCGTATTTCGTCTCGCTCAGGCGCAATCACGAATGAATAAC<br>GGTTTGGTTGATGCGAGTGATTTTGATGACGAGCGTAATGGCTGGCCTGTTGAAC<br>AAGTCTGGAAAGAAATGCACAAACTTTTGCCATTCTCACCGGATTCAGTCGTCACT<br>CATGGTGATTTCTCACTTGATAACCTTATTTTTGACGAGGGGAAATTAATAGGTTG<br>TATTGATGTTGGACGAGTCGGAATCGCAGACCGATACCAGGATCTTGCCATCCTAT<br>GGAACTGCCTCGGTGAGTTTTCTCCTTCATTACAGAAACGGCTTTTTCAAAAATAT<br>GGTATTGATAATCCTGATATGAATAAATTGCAGTTTCATTTGATGCTCGATGAGTT<br>TTTCTAAGTCGACATAACTTCGTATAATGTATGCTATACGAACGGTAGAAATTGCA<br>ATGCATCTGCCGAATGCCGTGTGGACGTAAGCGTGAACGTCAGGATCACGTTTCC<br>CCGACCCGCTGGCATGTCAACAATACGGGAGAACACCTGTACCGCCTCGTTCGCC<br>GCGCCACCATAAATCACCGCACCGTTCATCAGTACTTTCAGATAACACATCG |

**Supplementary video 1**

Depicts loss of the target plasmid encoding YFP in *E.coli* cells due to direct interference, related to Figure 2a,b. A single chamber of the microfluidic chip is shown. Phase contrast and fluorescent images were overlaid and compiled at 2-minute intervals starting from induction (0 h). Time is indicated in the bottom right in hours.

**Supplementary video 2**

Depicts loss of the target plasmid encoding YFP in *E.coli* cells due to priming, related to Figure 2e,f. A single chamber of the microfluidic chip is shown. Phase contrast and fluorescent images were overlaid and compiled at 2-minute intervals starting from induction (0 h). Time is indicated in the bottom right in hours.
