## Supplementary Methods for "Single Cell Variability of CRISPR-Cas Interference and Adaptation"

### 1 Master Equation description of the probability of plasmid loss

In order to test whether the distribution of the target clearance times by direct interference can be reproduced by a simple one-step process, we consider a model using a compound probability for binding of Cascade to the target and subsequent target removal from the system. In bacteria the number of targets is subject to maintenance which delays the removal of $M_0$ targets. For sake of simplicity we ignore this additional step, which has the advantage that the number of unknown parameters is kept to an absolute minimum. Because direct interference is a fast process, one can assume that target maintenance does not have a strong effect on the clearance time distribution. The Cascade number is not constant, but rather Cascade production is induced at the beginning of the experiment. This simplified model only depends on five parameters: the delay after induction for production of Cascade $\tau_c$, the Cascade production rate $\sigma$, the turn-over rate of Cascade $\lambda$, the number of targets per cell $M$, and the probability of a target removal event $p_d$. The number of targets in individual cells will be in general stochastic, however due to target maintenance one can assume that this distribution will be quite narrow. For this reason, we set $M_0 = 5$ [18].

The time dependent Cascade copy number is modelled as a production-degradation process with a delay $\tau_c$ and zero initial amount of Cascade: The bulk mean $\mu(t)$ is given by:

$$\mu(t) \quad = \quad \frac{\sigma}{\lambda}\theta(t-\tau_c)\left(1-e^{-\lambda(t-\tau_c)}\right).$$

By fitting this equation to Cascade concentration data for the bulk mean (Fig.4b of the main text), we estimate: $\tau_c = 34$ min, $\sigma = 3$ min$^{-1}$, and $\lambda = 0.0061$ min$^{-1}$ to obtain an average copy number of almost 500 Cascades per cell at steady state.

The removal of $M_0$ targets from the system is a First-Passage-Time problem. We formulate the simple Master Equation (ME) for the conditional probability $P_M(t)$ to find $M$ targets in a cell at a given time $t$:

$$\frac{dP_M(t)}{dt} \quad = \quad \mu(t)p_d(M+1)P_{M+1} - \mu(t)p_d M P_M,$$

where $p_d$ is the compound probability that within the time interval $\Delta t$ a Cascade molecule binds to a target and the target is subsequently removed from the system.

To obtain the First-Passage-Time distribution we need to determine the survival probability $S$ to find at least one target, which is simply given by $S = 1 - P_0$. $P_0$ is obtained by solving the above ME with the initial condition $P_M(t=0) = \delta_{MM_0}$:

$$P_0(t|M_0) \quad = \quad \left[1-e^{-p_d\int_0^t \mu(t')dt'}\right]^{M_0}.$$

$P_0(t|M_0) = 0$ for $t < \tau_c$ and because the state $M = 0$ is naturally an adsorbing boundary we readily find $\lim_{t\to\infty} P_0(t|M_0) = 1$. The First-Passage-Time distribution $FP_r(t|M_0)$ for target removal is given by the $FP_r = -dS/dt = dP_0/dt$:

$$FP_r(t|M_0) \quad = \quad M_0 p_d \mu(t)\left[1-e^{-p_d\int_0^t \mu(t')dt'}\right]^{M_0-1}.$$

Fitting this distribution to the empirical data (Fig. 2d of the main text) gives rise to $p_d = 4.4\times10^{-4}$ min$^{-1}$. The average target removal time $\tau$ is given by:

$$\tau \quad = \quad \int_0^\infty t' FP_r(t'|M_0)dt'.$$

Using the estimates for $p_d$, $\sigma$, $\lambda$, $\tau_c$, and $M_0 = 5$ we obtain $\tau \approx 94$ min. The fit of $FP_r$ to the data can be seen in Supplementary Methods Fig. 1a.

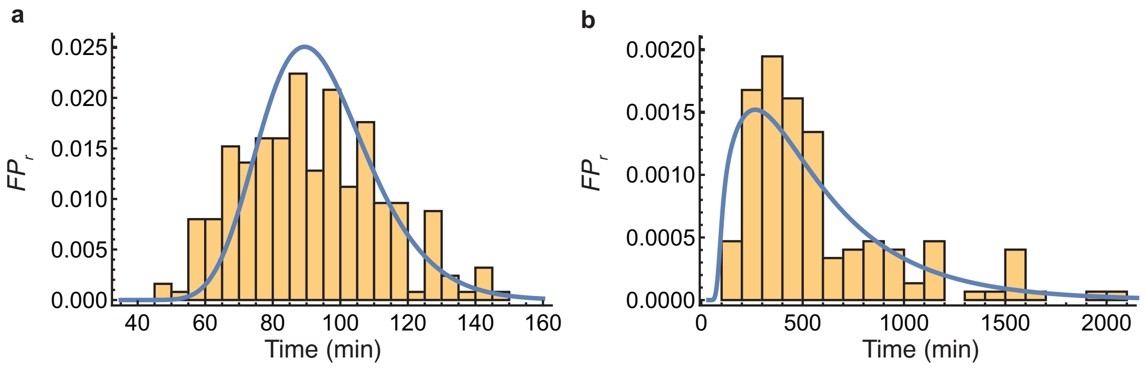

Supplementary Methods Figure 1: The fits of the simple one or two-step model to the data. **a**, fit of $FP_r$ (solid line) to the target removal time in the case of direct interference. **b**, fit of $FP_r$ (solid line) to the target removal time in the case of priming.

The simplified model yields a decent fit to the direct interference data. What about the target clearance during priming? To investigate whether this can be conceptually understood by a two-step process, first spacer acquisition subsequently followed by primed interference, we condition $FP_r$ on the time $\tau_p$ needed for spacer acquisition:

$$FP_r(t|M_0, \tau_p) \quad = \quad M_0 p_d \theta(t - \tau_p)\mu(t) \left[1 - e^{-p_d \int_{\tau_p}^{t} \mu(t')dt'}\right]^{M_0 - 1}.$$

The rationale behind this is that a Cascade molecule needs to bind to a target to produce the pre-spacers necessary for spacer acquisition before primed interference can happen. It follows $FP_r(t|M_0, \tau_p) = 0$ for $t < \tau_p$. Note that $\tau_p \geq \tau_d$, since in the absence of Cascade the probability of spacer acquisition is negligibly small. The distribution for $\tau_p$ is given by the First-Passage-Time distribution for the passage $M_0 \to M_0 - 1$: $FP_p = -dP_{M_0}/dt$:

$$FP_p(\tau_p|M_0) \quad = \quad M_0 p_p \mu(\tau_p) e^{-M_0 p_p \int_0^{\tau_p} \mu(t')dt'},$$

where $p_p$ is the compound probability that within the time interval $\Delta t$ one Cascade binds to a target, pre-spacers are produced and a spacer is integrated.

The distribution for the target removal times is given by:

$$FP_r(t|M_0) \quad = \quad \int_0^{\infty} FP_r(t|M_0 - 1, \tau_p) FP_p(\tau_p|M_0) d\tau_p.$$

The integral cannot be done analytically. Fitting $FP_r(t|M_0)$ to the experimentally obtained data for the distribution of target loss times during priming (Fig. 2e of the main text) yields $p_p = 10^{-6}$ min$^{-1}$. The fit of $FP_r$ to the data can be seen in Supplementary Methods Fig. 1b.

### 2 An agent-based model for stochastic biochemical kinetics of cell populations in microfluidic wells

Although a highly simplified description of our system, the results from the ME description show that the Cascade copy number is an important determinant in creating the variability in the PLT distribution in the case of direct interference. For priming, the distribution could be replicated by considering the process as the result of two subsequent steps, of which the spacer acquisition process creates the wide PLT distribution. However, this model of primed adaptation is highly simplified and does not give any mechanistic insight into the process of adaptation and interference in a growing cell population. To better understand how cell-to-cell variability and population dynamics affect CRISPR-Cas defense, we have developed a stochastic, agent-based simulation framework to analyse the kinetics of spacer acquisition and target loss. An agent-based approach allows us to keep track of the biochemical composition of individual cells in a growing population, as well as the inheritance of molecules and other cellular features in lineages. In this type of model, each cell is an agent, and there is no interaction between cells. For computational efficiency and to emulate the experimental set-up, the size of the cell population is kept constant. Results for this type of set-up, where the population size is constant, are identical as for a population experiencing exponential population growth, as long as the population size is sufficiently large (100-1000 cells) [17]. The intracellular reactions are governed by stochastic reaction kinetics which can be described by the Chemical Master Equation (CME). As an exact solution to the CME exists only for a handful of simple reaction networks, we use the stochastic simulation algorithm (SSA) [3], which provides trajectories which are consistent with the CME provided the rate constants are time-independent. When a reaction involves more than one molecular species, the propensity for this reaction to take place in some small time interval depends on the cellular volume. In our application, we are dealing with cells that are continuously in the exponential growth phase which violates the assumption of the SSA of constant propensities between reaction events. For this reason, we use the *Extrande* extension by Voliotis *et al.*, which allows us to efficiently simulate the reaction network containing time-dependent propensities [19].

#### 2.1 Model assumptions

Since the detailed mechanism of primed spacer acquisition in type I-E CRISPR-Cas systems is not yet completely known, we start out with a simplified model to see if this is sufficient to explain our data. Because primed adaptation is much more efficient than naive adaptation [14], we assume that the rate of naive adaptation is negligibly small over the time course of the experiment. The spacer composition of the CRISPR array is not modeled in detail. Rather, we assume that we start out with a crRNA sequence that matches the target, but is flanked by a non-consensus PAM. The effector complexes containing this spacer can still bind to the target DNA [10, 9], but with a binding affinity that is decreased up to a factor $100 - 150$ as compared to binding with a consensus PAM [5, 1]. Once the effector complex is bound to the target, Cas3-catalysed destruction of the target takes place [6]. Thus, the level of interference is associated with the level of effector complex binding [1].

Cas3-mediated destruction of targets is a source of substrates for spacer acquisition machinery, the Cas1-Cas2 complex, during primed adaptation [7, 11]. Intermediates of target DNA degradation are transient and quickly degrade after an initial burst. Abundant levels of Cas1 and Cas2 lead to robust spacer acquisition, by allowing Cas1-Cas2 to capture the transient intermediates of Cas3 action [11]. Since in our system Cas3, Cas1, and Cas2 are highly expressed, we assume the levels of these proteins are not rate-limiting within the scope of our model and thus do not explicitly model their abundances. Furthermore, in agreement with previously published work, we assume cells have a target maintenance system that is controlled by logistic dynamics in order to keep the target concentration at its target level [12]. In addition, targets and target-containing configurations are actively partitioned between daughter cells [8, 13] according to a multi-hypergeometric distribution, with each daughter receiving on average half of the mother cell's targets. All other proteins are partitioned according to a Binomial distribution, where the ratio of daughter cell sizes determines the probability of each molecule ending up in one of two daughter cells. We model synthesis of CRISPR proteins as a Poisson process, in which proteins are produced in geometrically distributed bursts to capture the effect of transcriptional bursting [4]. We assume all molecular species are stable on the timescale of the experiment (i.e. no degradation), with the exception of the free crRNAs (not loaded in Cascade) and the DNA fragments that are the result of interference, which have a short lifetime.

#### 2.2 Algorithm outline

For the agent-based model, we have adapted the First-Division Algorithm by Thomas [16] to include the *Extrande* extension to the SSA. Furthermore, we keep the population size constant by randomly selecting a cell to be removed from the population in the event of a cell division. The steps to replicate our experimental set-up are described below.

1. **Population initialization:** At time $t = 0$, initialize $N$ cells by assigning to each cell an age $t_i \sim \mathrm{U}(-\log(2)/\mu_p, \log(2)/\mu_p)$, a growth rate $\mu_i \sim \mathrm{Lognormal}(\mu_p, \sigma_p^2)$ and molecule count $x_i$. Select division size $V_{d,i} \sim \mathrm{Lognormal}(\mu_{V_D}, \sigma_{V_D}^2)$ and compute generation time $t_{gen,i}$ as $\log(V_{d,i}/V_{b,i})/\mu_i$, where $V_{b,i}$ is the birth size. This determines the division time of the cell which is defined as $t_{d,i} = t_i + t_{gen,i}$.

2. **Biochemical reactions:** Determine the next dividing cell: $j = \mathrm{argmin}_i(t_{d,i} - t_i)$. Determine $\Delta t$ from $\min(t_{d,j} - t_j, L)$, where $L$ is *Extrande*'s look-ahead horizon. Advance the molecule numbers of each cell independently from age $t_i$ to $t_i + \Delta t$ using the *Extrande* algorithm and advance time from $t$ to $t + \Delta t$.

3. **Cell division:** When $t = t_{d,j}$, replace the dividing cell by two newborn daughter cells of zero age. The birth size of both daughters is determined as $V_{b,D_1} = \mathrm{Normal}(\mu_{V_B}, \sigma_{V_B})V_{d,j}$ and $V_{b,D_2} = V_{d,j} - V_{b,D_2}$. Assign to one of these a molecule number distributed according to the Binomial distribution (proteins) and the Multi-hypergeometric distribution (targets and target configurations), depending on the mother's molecule count $x_j$ and the daughter's size ratio to the mother cell $\frac{V_{b,D_1}}{V_{d,j}}$, and assign the remaining molecules to the other daughter. Assign to each daughter independently a growth rate $\mu_i$, division volume $V_{d,i}$, and compute corresponding division time. To ensure a constant population size, randomly select a cell to be deleted from the population.

4. **Repeat:** Repeat from 2. until $t = t_{\mathrm{final}}$.

#### 2.3 Molecular mechanism and model parameters

Each cell in the population contains a pool of biochemical species that can interact with each other through biochemical reactions, as described in step 2. We distinguish between the targets $P$, the CRISPR array $A$, which codes for a spacer $crRNA$ matching a sequence on the target, and the surveillance protein $Cascade$. Together with the crRNA, the Cascade protein makes up the effector complex $E$. When the effector complex encounters a target it can bind, albeit with a low affinity in the case of a non-consensus PAM on the target, forming a complex $EP$. Destruction of the target can then take place, producing DNA fragments $F$. One of these fragments can be integrated into the CRISPR array $A$ as a new spacer, transforming the array to $A^*$ which can now also express the newly acquired crRNA, $crRNA^*$, in addition to the spacer that was already present. The effector complex containing the new spacer, $E^*$ has a higher binding affinity for the target. These biochemical reactions are governed by the equations described in Table 1.

The size of individual cells increases exponentially with a constant elongation rate throughout the cell cycle. Cellular length is used as a measure for cell size, as *E. coli* cell width remains approximately constant throughout the cell cycle and thus the cellular volume is linearly proportional to the cell length [15]. Growth parameters were chosen to be representative for our experimental data. As no kinetic data are available on individual reactions of the adaptation and interference processes, these parameters were calibrated to qualitatively agree with the experimentally determined target loss time distributions from the direct interference and priming conditions and previously published abundances of *cas* abundances [2]. Unless stated otherwise, the growth parameters used were $\mu_p = \log(2)/70$, $\sigma_p = 0.2$, $\mu_{V_B} = 0.5$, $\sigma_{V_B} = 0.07 \cdot \mu_{V_B}$, $\mu_{V_D} = 3.9$, $\sigma_{V_D} = 0.11 \cdot \mu_{V_B}$, $p^* = 5$. To simulate the direct interference condition with the same model, we simply modify the initial state of the system such that the spacer array consists of crRNA$^*$, which is flanked by the consensus PAM sequence.

#### 2.4 Cascade variability impacts the probability of spacer acquisition

In the main text of the manuscript we have shown that in priming, increased variability in the expression of Cascade can lead to faster spacer acquisition on average (Fig. 5c of manuscript). In simulations of the agent-based model, variability of the Cascade protein concentration is controlled through the protein production rate $k_1$ in coordination with the average protein burst size $b_c$: to modify Cascade variability while maintaining a constant concentration, $b_c$ is multiplied by a factor $a$ while $k_1$ is multiplied by its inverse, $\frac{1}{a}$. In Fig. 5, $a = 100$ which leads to an increase of the coefficient of variation of the Cascade concentration at steady state from $CV = 0.02$ (low Cascade variability) to $CV = 0.42$ (high Cascade variability).

We will now illustrate how higher Cascade variability can lead to faster spacer acquisition by considering two scenarios, and comparing the cumulative probability of the time until spacer acquisition for the simplified two-step model, which is given by

$$FP_{SA}(t|M_0) = 1 - e^{-M_0 p_p \int_0^t \mu(t')dt'}.$$

First, we consider a cell which has a constant Cascade level of 500 copies at any point in time between $t = 0 - 1000$ min, and plot the corresponding cumulative spacer acquisition probability (Supplementary Methods Fig. 2a). Second, we consider a second cell in which Cascade is not constant but rather appears as a shorter 'burst' of 2500 copies from $t = 200$ min until $t = 400$ min, and 0 copies at any other time (Supplementary Methods Fig. 2a). The cumulative spacer acquisition probability for the second cell reaches 1 faster than for the first cell (Supplementary Methods Fig. 2b), despite the two cells having the same average Cascade concentration over the course of 1000 minutes. This suggests that the effects of upwards fluctuations can outweigh the downward fluctuations.

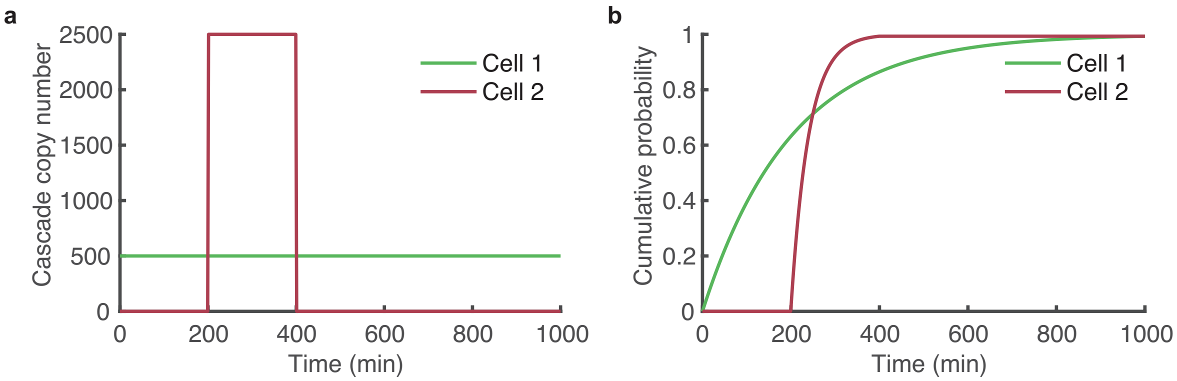

Supplementary Methods Figure 2: **a**, Cascade copy number of two cells with the same average over time. Cell 1 has a constant copy number of 500 Cascades, while in cell 2 Cascade is present transiently at 2500 copies between 200 and 400 minutes. **b**, Cumulative probability of time until spacer acquisition for the two cells with Cascade copy numbers as described in panel a ($M_0 = 1$, $p_p = 0.00001$).

| Phase | Reactions |
|---|---|
| target replication | $P \xrightarrow{k_0/(1+((P/V_t)/p_0)^2)} 2P,$<br>$p_0 = \frac{p^*}{V_t}/\sqrt{(\frac{k_0}{\mu} - 1)}$ |
| Expression | $G \xrightarrow{k_1(t)} G + b_P \cdot Cascade$<br>$k_1(t) = \frac{k_1}{1+exp(-k_d t)}$<br><br>Before spacer integration<br>$A \xrightarrow{k_2} A + b_c \cdot crRNA$<br><br>After spacer integration<br>$A^* \xrightarrow{k_2} A^* + b_c \cdot crRNA + b_c \cdot crRNA^*$<br><br>$crRNA + Cascade \xrightarrow{k_3} E$<br>$crRNA^* + Cascade \xrightarrow{k_3} E^*$ |
| Interference | $E + P \underset{k_5}{\overset{k_4}{\rightleftharpoons}} EP$<br>$E^* + P \underset{k_7}{\overset{k_6}{\rightleftharpoons}} EP^*$<br>$EP \xrightarrow{k_8} E + b_F \cdot F$<br>$EP^* \xrightarrow{k_8} E^* + b_F \cdot F$<br>$F \xrightarrow{k_9} \emptyset$ |
| Primed adaptation | $F + A \xrightarrow{k_{10}} A^*$ |

Supplementary Methods Table 1: Overview of the reactions in the model for primed adaptation.

| **Reaction** | **Parameter** | **Value** $[min^{-1}]$ |
|---|---|---|
| target replication | $k_0$ | 0.125 |
| *Cascade* production | $k_1$ | 2.4 |
| $crRNA/crRNA^*$ transcription | $k_2$ | 10 |
| $crRNA/crRNA^*$ degradation | $k_3$ | 0.014 |
| $crRNA-Cas/crRNA^*-Cas$ effector complex formation | $k_3$ | 0.01 |
| $E-P$ binding affinity | $k_4$ | $1e^{-5}$ |
| $EP$ dissociation | $k_5$ | $1e^{-4}$ |
| $E^*-P$ binding affinity | $k_6$ | $1e^{-3}$ |
| $EP^*$ dissociation | $k_7$ | $1e^{-4}$ |
| target degradation | $k_8$ | 1 |
| Fragment degradation | $k_9$ | 1 |
| Spacer integration | $k_{10}$ | 0.25 |
| *Cascade* burst size | $b_P$ | 3 |
| $crRNA/crRNA^*$ burst size | $b_c$ | 3 |
| DNA fragment burst size | $b_F$ | 5 |
| Post-induction delay of protein production | $k_d$ | 0.025 |

Supplementary Methods Table 2: Reaction rates used in simulations
